# Plasma cells are formed in waning germinal centers via an affinity-independent process

**DOI:** 10.1101/2023.04.13.536825

**Authors:** Henry J. Sutton, Brian J. Parker, Mariah Lofgren, Cherrelle Dacon, Deepyan Chatterjee, Xin Gao, Hannah G. Kelly, Robert A. Seder, Joshua Tan, Azza H. Idris, Teresa Neeman, Ian A. Cockburn

## Abstract

Long-lived plasma cells (PCs) secrete antibodies that can provide sustained immunity against infection. It has been proposed that high affinity cells are preferentially selected into this compartment, potentiating the immune response. Here we used single cell RNA-seq to track the development of Igh^g2A10^ cells, specific for the *Plasmodium falciparum* circumsporozoite protein (*Pf*CSP) within the germinal center (GC). We identified cells differentiating into memory B cells (MBC) or PCs and estimated their affinity by V(D)J sequencing. While pre-memory cells were of lower affinity than GC B cells generally, the affinity of pre-PC cells was indistinguishable. Rather, a larger proportion of cells differentiated into PCs in waning GCs. These later emigrants replaced early arrivals in the bone marrow allowing the development of a high affinity PC compartment.

## Introduction

Following vaccination, antibody responses to many pathogens can be sustained for periods of decades, in some cases conferring life-long protection against disease ^1,2^. Sustained antibody production is maintained by long-lived Plasma Cells (PCs) which reside in multiple organs, but notably the bone marrow (BM) and spleen ^3,4^. BM PCs secrete high affinity antibodies that have undergone a process of somatic hypermutation and selection called affinity maturation ^5,6^. This process takes place in structures called germinal centers (GC) where antigen specific B cells cycle between the dark zone (DZ), the primary site of cell proliferation and the light zone (LZ) where B cells encounter cognate antigen and obtain help from T cell ^7-9^. Cells that fail to bind antigen and obtain T cell help will die facilitating a Darwinian selection process called affinity maturation. Cells that survive make a fate choice between continuing to cycle in the GC or exiting as a Memory B cell (MBC) or PC.

It has long been noticed that MBCs are generally of lower affinity than PCs ^10^. It has also been shown that low affinity cells in the GC are preferentially selected into the MBC compartment ^11,12^. PC formation requires T cell help and it is proposed that high affinity B cells obtain more antigen for presentation on MHC, facilitating their differentiation into antibody secreting cells ^13-15^.

However, paradoxically, access to stronger T cell help by high affinity cells also aids their recycling into the DZ ^16^. Another model based on BrdU pulse-chase labelling of B cells proposes that longer lived PCs in the BM are preferentially formed in the late GC ^17^. As late GCs contain more affinity matured cells, this temporal switch model would not require the preferential differentiation of high affinity GC cells into PCs. A recent study using tamoxifen lineage tracking mice challenged the temporal switch model by showing that PCs emerge at a constant rate from the GC ^18^. However similar lineage tracing experiments have proposed that early GC emigrants are replaced by later emigrants, explaining why most long lived PCs derive from the late GC ^19,20^. To examine the role of affinity in this process we tracked Igh^g2A10^ cells specific for the malaria vaccine antigen *P. falciparum* circumsporozoite protein (*Pf*CSP) by single cell RNA-seq in the GC. Transcriptomic data allowed us to identify Pre-Memory cells (PreMem) and Pre-PCs (PrePC) among LZ B cells, while BCR sequencing allowed us to estimate the affinity of these cells and thus understand the relationship between affinity and cell fate choices in the GC.

## Results

### Stochastic cell fate choice and stereotypic affinity maturation in Igh^g2A10^ cells

Igh^g2A10^ mice have the unmutated common ancestor (UCA) of the heavy chain of the 2A10 mAb, specific for the *Pf*CSP repeat domain knocked into the IgM locus ^21^. Because the Igh^g2A10^ heavy chain is free to pair with any light chain we reasoned we might detect populations in this mouse that vary in their ability to bind to *Pf*CSP. In agreement with this we identified populations binding high, intermediate, and low amounts of *Pf*CSP probe relative to their IgM expression (Figure 1A). Single cell BCR sequencing revealed that all *Pf*CSP-binding cells utilized the *Igkv10-94* gene, which is also used by the 2A10 mAb ^22,23^. However, in the different *Pf*CSP-binding populations this *Igkv* gene was paired with *Igkj2* (J2; high), *Igkj2* (J1; intermediate) or *Igkj4* (J4; low) genes resulting in different aromatic amino acids being present at position LC_116 (Figure 1B). ELISA of recombinant antibodies (rAbs) carrying each light chain confirmed that these changes resulted in expected reductions in affinity compared to the 2A10 UCA (Figure 1C).

**Figure 1.**
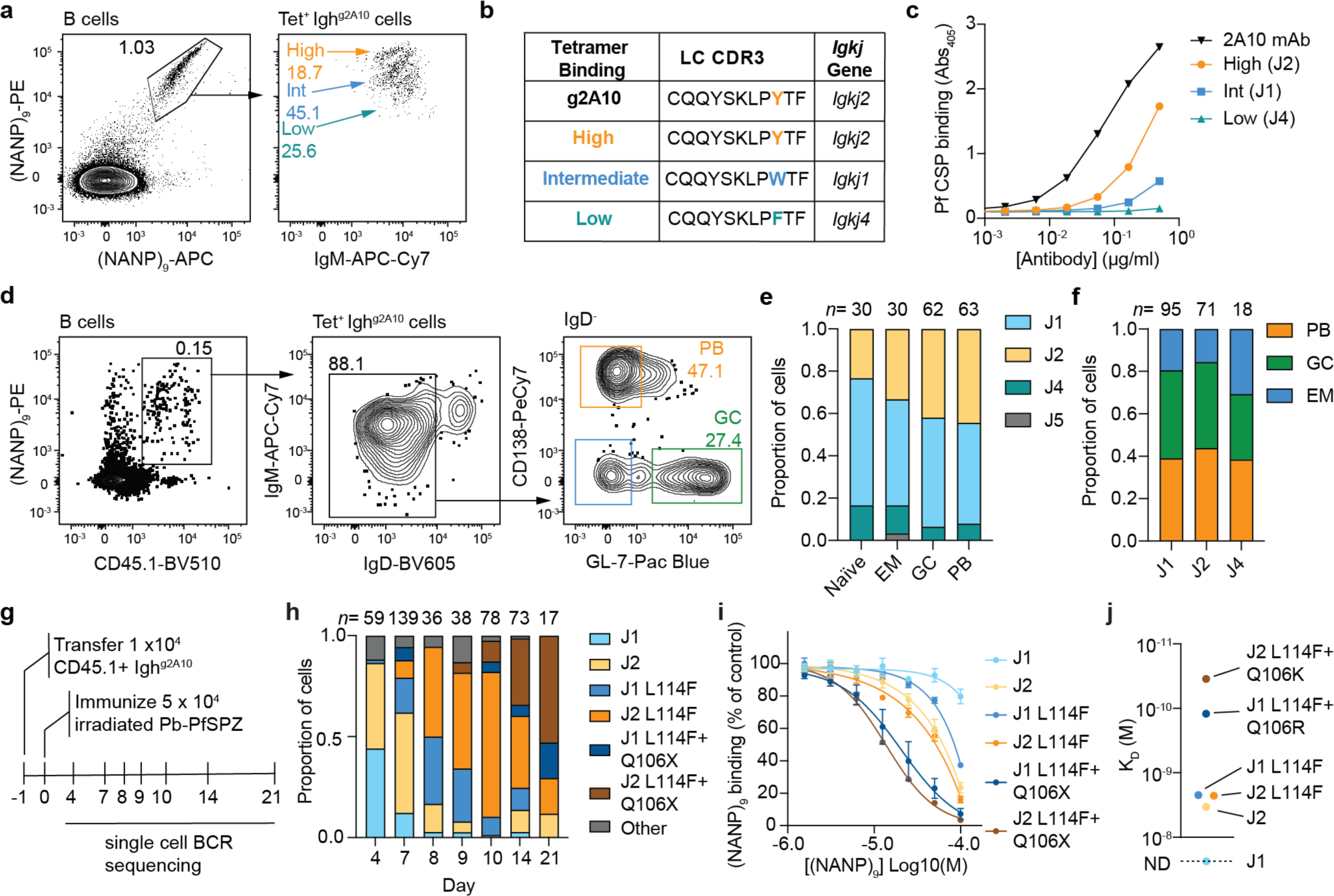
Affinity determinants and their influence on early cell fate choice in Igh^g2A10^ cells. **A**. Flow cytometry plots demonstrating 3 distinct populations of Igh^g2A10^ B cells separated by their binding to P*f*CSP repeat ((NANP)_9_-PE and (NANP)_9_-APC probes **B**. The LC-CDR3 amino acid sequence and *Igkj* gene used by each probe binding population compared to the predicated germline 2A10 LC-CDR3. **C**. ELISA showing the P*f*CSP binding ability of recombinant antibodies generated using sequences from (C) compared to 2A10 **D**. Representative flow plots demonstrating the gating strategy used to sort *Pf*CSP-specific Igh^g2A10^ PB, GC and EM B cells 4 days post PbPf-SPZ immunizations. **E**. *Igkj* gene usage in either naïve Igh^g2A10^ B cells or PB, GC and EM B cells 4 days post immunization with PbPf-SPZ **F**. Proportions of Igh^g2A10^ PB, GC or EM B cells utilizing either the J1, J2 or J4 *Igkj* gene **G**. Experimental schematic of time course analysis of Ighg2A10 GC evolution. **H**. Frequency of J gene usage and occurrence of L114X and/or Q106X mutations 4, 7, 8, 9, 10,14 and 21 days post immunization with PbPf-SPZ **I**. Competition ELISA showing the binding of the indicated to rAbs to plate bound (NANP)_9_ peptide in the presence of the indicated concentrations of soluble (NANP)_9_ peptide; Mean ± SEM shown from 3 independent experiments. **J**. Affinities (KD) of mAbs carrying the indicated variants; all values represent the geometric mean of binding signals collected in duplicate from n = 2 independent experiments. ND, could not be determined.

It has been proposed that high affinity cells preferentially become PCs, while GCs are formed from cells with lower affinity, though others have suggested this initial fate choice is stochastic ^24-26^. To test this hypothesis Igh^g2A10^ cells were transferred to mice which were subsequently immunized with irradiated *P. berghei* sporozoites expressing *Pf*CSP in place of their endogenous *P. berghei* CSP molecule (PbPf-SPZ) ^27^. Immunization with irradiated sporozoites is an established method for generating sterilising immunity against malaria in mice and humans ^28,29^. Four days after immunization, plasmablasts (PB), GC B cells, and early memory (EM) B cells were single cell sorted and the light chains sequenced (Figure 1D). Overall, when cells were separated by cell fate (χ^2^=11.3; df=9; p=0.25) or J chain use (χ^2^=1.89; df=4; p=0.76) there was no clear difference in the propensity to enter a particular cell fate (Figure 1E-F). Thus, our data are consistent with early cell fate choices being largely stochastic.

To understand how Igh^g2A10^ cells mature in the GC we extended the earlier analysis to multiple time points after immunization (Figure 1G). Our previous investigation of the 2A10 antibody showed that most affinity maturation occurs in the LC so we focussed on this chain ^22^. At day 7 some cells carried a C328T mutation that codes for a LC_L114F substitution (Figure S1A-B; Figure 1H). This mutation was more common initially in J1 cells which had become rarer than at day 4, perhaps because of greater selective pressure on these initially lower affinity cells (Figure 1H). By day 8 the LC_L114F mutation had swept through the GC and was found in ∼80% of cells (Figure 1H). By day 21 an LC_Q106X substitution occurring in ∼50% of GC B cells had become next most common (Figure 1H). At this position multiple different changes were observed, notably LC_Q106R on the J1 light chain (15/30 mutated J1 chains), with LC_Q106K (22/75) or LC_Q106L (18/75) on the J2 light chain (Figure S1A-B; Figure 1H). Competition ELISA of rAbs carrying these mutations confirmed that these mutations resulted in progressively higher affinity for the (NANP)_9_ repeat (Figure 1I). To precisely measure the effect of these mutations on BCR affinity we generated antigen-binding fragments (Fabs) containing these mutations and measured binding to *Pf*CSP by surface plasmon resonance (SPR; Figure S1C). The SPR analysis revealed that the L114F mutation on its own resulted in a modest (1.49-fold) increase in affinity over the J2 UCA. Improvements in affinity due to the L114F substitution could not be determined in the J1 lineage due to the negligible binding signals of the J1 UCA. Interestingly the C328T mutation occurs at a WG<u>C</u>W super-hotspot for AID targeting ^30^, potentially explaining the rapid selective sweep of this mutation despite a relatively small affinity advantage. Additional LC_Q106K/Q106R mutations substantially enhanced affinity in both lineages, conferring a 65-fold increase in affinity to the J2/L114F light chain and an 18.2-fold increase in affinity to the J1/L114F light chain respectively (Figure 1J).

### Single cell RNA sequencing allows the identification of PrePCs and PreMems within the GC

Because we can see stereotypic selective sweeps occurring among GC B cells based on Igh^g2A10^ cells we exploited this feature to determine if these affinity changes were related to cell fate decisions in the GC. Specifically, we aimed to use single cell RNA-seq to identify PrePCs or PreMem cells among Igh^g2A10^ cells in the GCs and combine this transcriptomic data with V(D)J sequencing to determine if these PreMem and PrePC populations contained generally higher or lower affinity cells (Figure 2A). Accordingly, we sorted and sequenced IgD^-^ Igh^g2A10^ cells from 5 mice each at days 7 and 21 post immunization with PbPf-SPZ. Days 7 and 21 were chosen as at each of these days ∼50% of GC B cells carried the LC_L114F and LC_Q106X mutations respectively. To facilitate the identification of PrePC and PreMem populations we included CITE-seq ^31^ Abs specific for the PrePC marker CD69 and CD38 which is upregulated on PreMem cells ^15,32,33^. We also included antibodies specific for CD11c and CXCR3 to distinguish atypical B cells from conventional MBCs ^34,35^.

**Figure 2:**
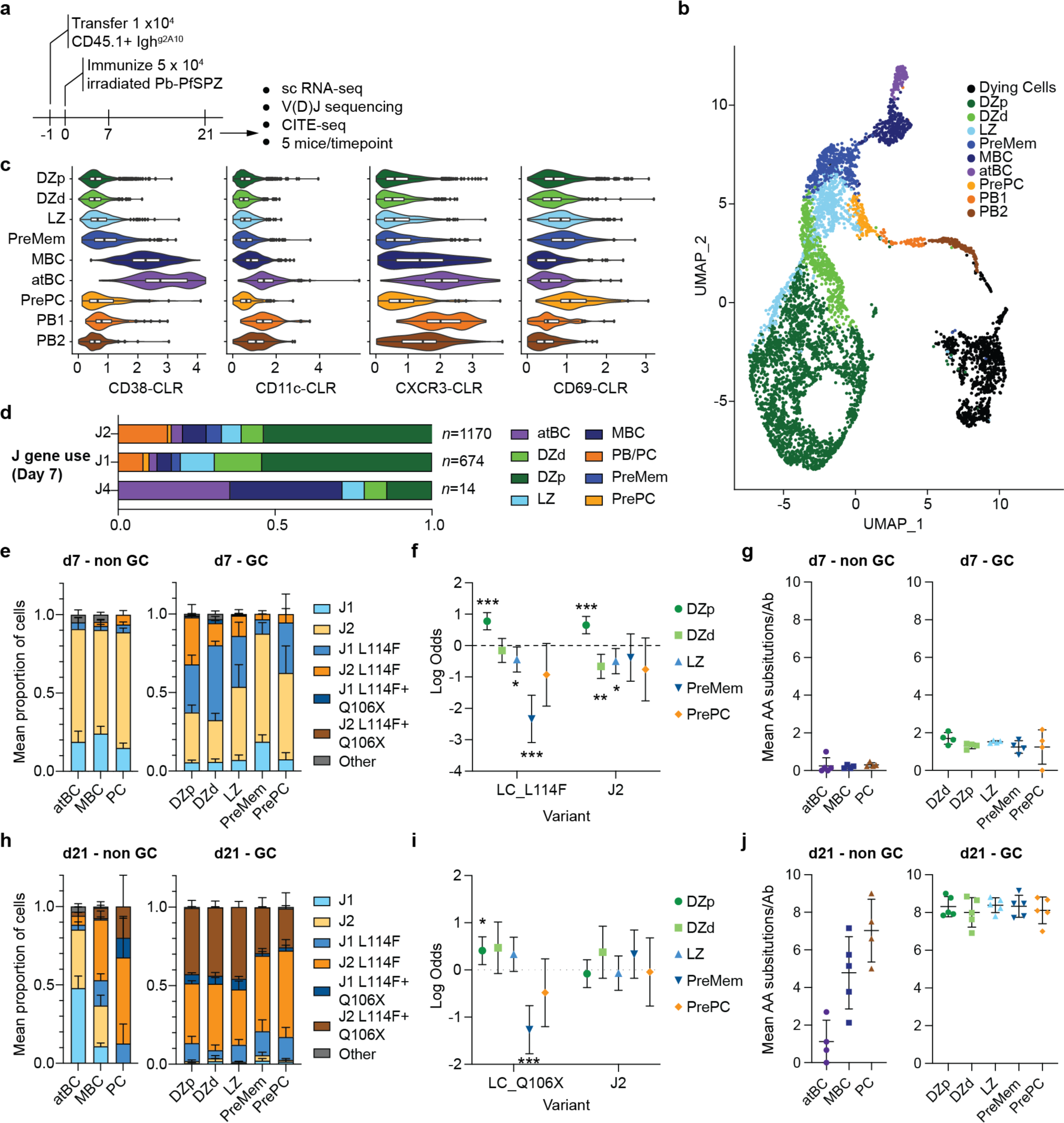
Single cell RNAseq identifies PreMem and PrePC populations within the GC. **A**. Experimental Schematic. **B**. Unsupervised clustering of P*f*CSP -specific Igh^g2A10^ B cells pooled from all mice visualized using UMAP. Each cell is represented by a point and coloured by cluster **C**. Violin and box plots showing the centred log ratio (CLR)-normalized expression of surface proteins used to identify MBC, PreMem and PrePC populations using CITE-seq. D. Proportion of J1, J2 and J4 cells in each B cell population identified by single cell transcriptomics 7 days post immunization. **E**. Frequency of J gene usage and occurrence of LC_L114F and LC_Q106X mutations 7 days post immunization in non-GC and GC populations; data are expressed as mean ± SD of the 5 mice. **F**. Likelihood of the LC_L114F mutation being present in different GC populations 7 days post immunization; data are expressed as Log Odds ±95% CI from logistic regression analysis controlling for mouse as random effect. **G**. Mean AA substitutions in each mouse 7 days post immunization in non-GC and GC populations; data are expressed as mean ± SD. **H**. Frequency of J gene usage and occurrence of L114X and/or Q106X mutations 21 days post immunization in non-GC and GC populations; data are expressed as mean ± SD of the 5 mice. **I**. Likelihood of the LC_Q106X mutation being present in different GC populations 21 days post immunization; data are expressed as Log Odds ±95% CI from logistic regression analysis controlling for mouse as random effect. **J**. Mean AA substitutions in each mouse 21 days post immunization in non-GC and GC populations; data are expressed as mean ± SD.

Dimensionality reduction analysis showed 4 major populations of cells (Figure 2B) which we identified as dying cells, GC B cells, PCs and MBCs using a combination of gene set enrichment and the expression of hallmark genes (Figure S2A-B). The dying cells had higher levels of mtRNA (Figure S2A) and might include cells harmed by experimental manipulation as well as those counter selected in the GC and so were excluded from downstream analysis. Among MBCs, we could detect a population of cells with high surface CD11c expression that were identified at atBCs (Figure 2C). The PC population could be divided into PC1 and PC2 based principally on the expression of cell cycle genes (Figure S2C). Among GC B cells we identified two clusters of DZ cells based on their cell cycle stage and the expression of *Aicda, Mki67* and *Ccnb1* (Figure S2B-C) which conform to a previous classification of DZ cells as proliferating (DZp) or differentiating (DZd) ^36^. Three populations were observed with high expression of LZ genes including *Cxcr5, Cd86* and *Cd83* (Figure 2B and Figure S2B). Cells from the first of these clusters were simply defined as LZ B cells, however, a second cluster was located proximal to the MBC population in our UMAP projection (Figure 2B) and had a similar gene expression profile to MBCs (Figure S2A). CITE-seq analysis revealed that these had slightly higher surface expression of CD38 so this population was designated PreMem (Figure 2C). The third LZ cluster, similar to one identified in a recent study ^37^, was enriched for genes including *Cd86, Irf4, Myc, Cd40* and *Icam1* which have been associated with a population of Bcl6^low^ CD69^hi^ PrePC cells (Figure S2A-C). These cells also had increased surface expression of CD69 (Figure 2C) further supporting their designation and PrePC cells ^15^. Pseudotime analysis showed that when LZ B cells were set as the root, 3 distinct trajectories could be seen, one leading to DZ recycling and the other two passing through either PreMem to MBC or PrePC to PC (Figure S2D).

### PC differentiation is affinity independent

Having identified PreMem and PrePCs we asked if we could relate these cell fates to the presence of affinity-increasing mutations. 7 days after immunization, ∼63% were J2, ∼36% were J1 while <1% were J4. While the cellular makeup of the of J1 and J2 populations were similar at day 7, J4 cells were largely found in the memory compartment (Figure 2D), suggesting that even if these cells were capable of entering the GC (Figure 1E-F) they were rapidly outcompeted. atMBC, MBC and PC cells were mostly unmutated, however ∼50% of cells in the GC carried the LC_L114F mutation, though this was more common in the DZd, DZp, and LZ populations compared to the PreMem and PrePC populations (Figure 2E). Logistic regression analyses revealed that PreMem cells were much less likely to contain the LC_L114F mutation compared with other cell fates (OR = 0.096, 95% CI [0.046,0.21]). Conversely DZp cells were enriched relative to other cells in the LC_L114F mutation (OR=2.7, 95% CI [1.66,2.86] and *Igkj2* gene (OR = 1.92, 95% CI = [1.66,2.86]) consistent with rapid selection of this mutant at this time (Figure 2F). One possibility is that PreMem are lower affinity because they began to differentiate before the LC_L114F selective sweep had begun. However, overall levels of mutation were similar in this population compared to other GC populations, suggesting these cells had only just exited GC cycling (Figure 2G).

By day 21 the LC_L114F mutation had swept through all GC populations, and the small number of spleen PC cells; however, some MBC and most atBC were unmutated even at day 21 suggesting these cells are either GC-independent or formed very early in the response (Figure 2H). Because the LC_L114F mutation was ubiquitous in the GC and no longer useful for distinguishing affinity, we examined the prevalence of mutations at LC_Q106. These were present in roughly 50% of DZd, DZp and LZ cells, but only around 30% of PreMem and PrePC cells (Figure 2I). Logistic regression analysis revealed that the LC_Q106X mutations were under-represented in PreMem cells (OR = 0.28, 95% CI [0.17-0.47]), and over-represented in DZp cells (OR = 1.50, 95% CI [1.12,2.02]) (Figure 2I). There was no evidence that cells with LC_Q106X mutations were preferentially selected into the PrePC compartment (OR = 0.61, 95% CI [0.30,1.27]). Finally, as at day 7, overall mutation frequencies were similar among all GC populations in all mice indicating recent differentiation into the PrePC or PreMem compartments (Figure 2J).

### Lineage analysis of Igh^g2A10^ GC B cells

We were concerned that our results were biased by focussing on a small number of pre-selected mutations. We therefore examined the frequency of other mutations in the Igh^g2A10^ lineage and their associations with cell fate. After LC_114 and LC_106, mutations at amino acids HC_39, HC_59 and HC_68 were the next most common across all mice (Figure 3A; Figure S3A-B). Collectively mutations at the top 10 positions accounted for around 40% of the mutation burden Igh^g2A10^ cells (Figure 3B). Phylogenetic analysis of cell lineages at day 21 revealed that these mutations typically arose close to the root of the trees and were responsible for small selective sweeps among the GC B cells (Figure 3C-D; Supplementary dataset 3). These mutations generally arose independently of each other, though mutations at positions HC_39, HC_40 and the light chain CDR3 were slightly favoured when a LC_Q106X mutation was present (Figure S3C).

**Figure 3:**
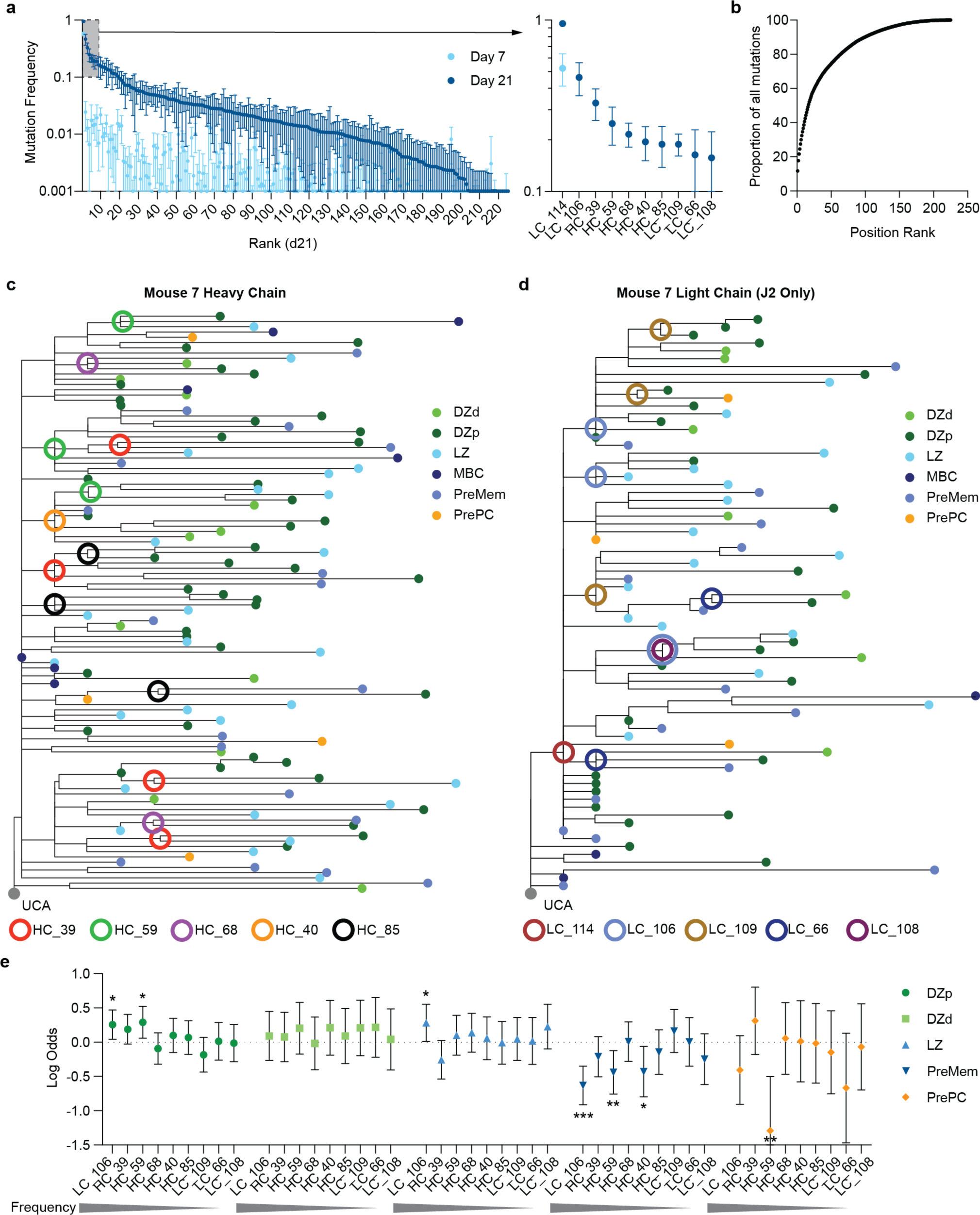
Mutational dynamics in the germinal center. Mice were immunized and analyzed as in Figure 2A. **A**. Frequency (mean ± SEM) of mutation at each HC and LC AA position in each mouse ranked from highest to lowest, inset shows the frequency of the top 10 mutations at day 21 **B**. Proportion of mutations accounted for by mutations at each position, ranked from most to least common. **C**. and **D**. Phylogenetic trees based on heavy chain (**C**) and light chain (**D**) mutations linking the Igh^g2A10^ cells in a representative mouse (mouse 7); closed circles at the tips represent the phenotype of the cell, open circles represent the nodes where common mutations arise. **E**. Likelihood of each of the top 10 most common mutations being found in cells in different GC populations; data are expressed as Log Odds ±95% CI from main effect logistic regression analysis controlling for mouse as random effect.

To address whether any specific mutations were associated with cell fate we performed a main-effects logistic regression analysis which showed that mutations at 3 of the 5 most mutated positions (LC_106; HC_59; HC_40) were significantly under-represented in PreMem cells (Figure 3E). Notably the HC_N59I mutation, has been shown to contribute to the affinity of the 2A10 mAb^22^. Again, no clear signature could be determined among PrePC cells though mutations at HC_59 were significantly rarer in this population. Overall, these data support the hypothesis that PreMem cells are generally derived from lower affinity cells in the GC but that PrePCs differentiate in an affinity independent manner.

### Late GC emigrants replace early emigrants in the BM

The lack of a clear affinity signature for PrePCs led us to examine the hypothesis that PrePCs preferentially emerge late in the GC reaction ^17^. Comparison of the B cell populations at days 7 and 21 showed that DZ populations made up a smaller proportion of GC B cells while the proportion of PrePCs had increased from ∼1% to 3% (Figure S4A, Figure 4A). To investigate the kinetics of GC exit we performed flow cytometry analysis of the GC and PrePC response from mice that had received Igh^g2A10^ cells and been immunized with PbPf-SPZ (Figure 4B). Based on our single cell RNA seq analysis PrePCs were identified as cells with elevated expression of Irf4, CD86 and CD69, however while cells that upregulated all these markers could be detected (Figure S4B), these cells did not form a distinct population that could be objectively gated. Accordingly, to estimate the number of PrePCs we quantified the number of GC B cells that were above the 80^th^ percentile for all three markers (Figure S4B). This revealed an excess of Irf4^hi^ CD69^hi^ CD86^hi^ cells compared to the proportion (0.8%) that would be expected by chance (Figure 4C). Consistent with later GCs favouring PC formation the estimated proportion of PrePC cells increased over the course of the GC reaction, but in terms of absolute numbers this still corresponded to a peak output of PrePCs at day 10 when the GC is largest but with a slow decline while the total GC wanes rapidly (Figure 4D). This relatively flat output of PrePC is in broad agreement with a recent study examining PC formation using Blimp1 fate-tracking models ^18^.

**Figure 4:**
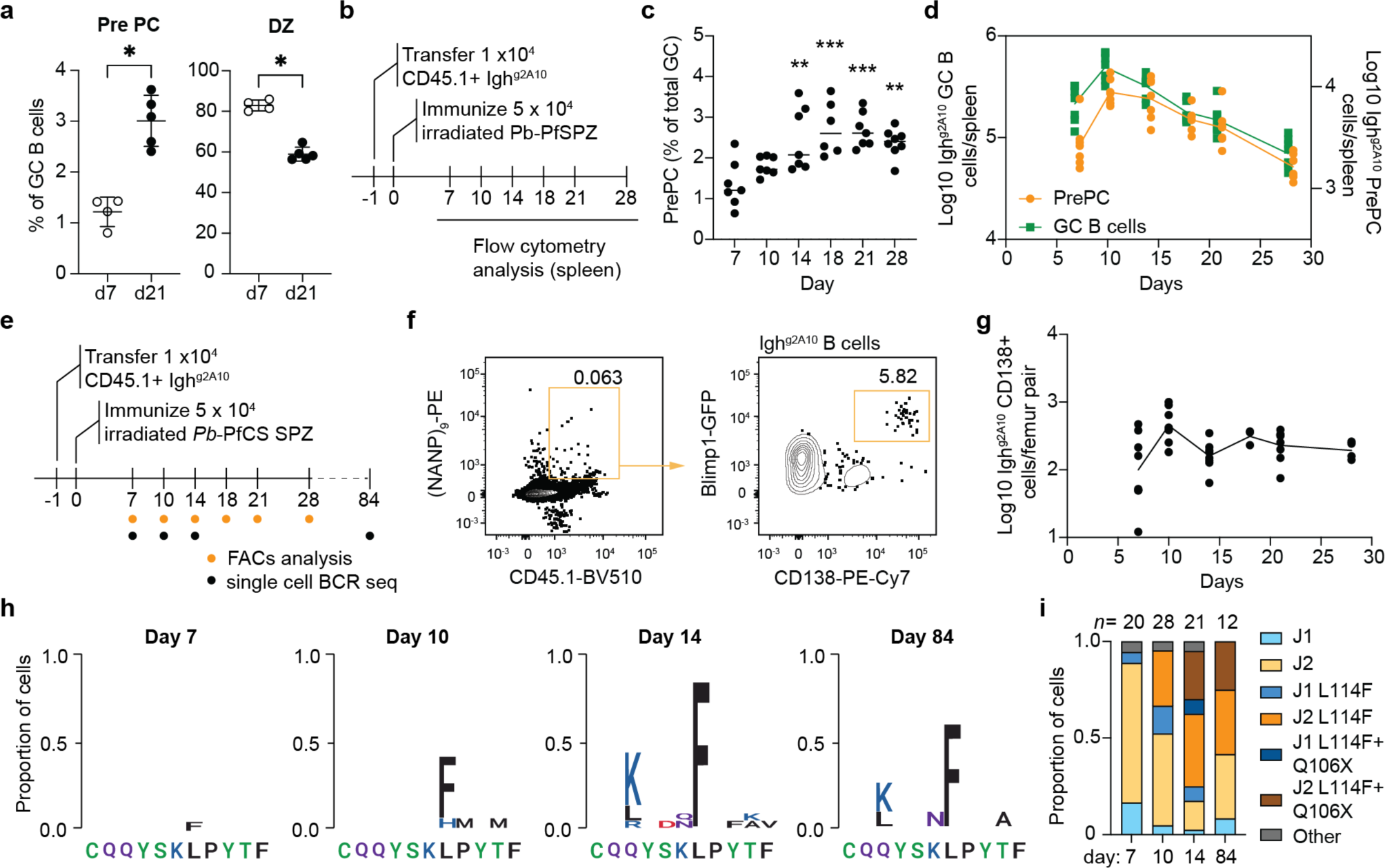
PC formation is favoured in waning GCs. **A**. Percentage of *Pf*CSP-specific Igh^g2A10^ GC B cells that are PrePCs or DZ cells at 7 and 21 days post immunization as identified by single cell RNA-seq as described in Figure 2A **B**. Experimental Schematic to examine PrePC populations in the spleen. **C**. Estimated PrePCs as percentage of total GCs over time. **D**. The total number of *Pf*CSP-specific Igh^g2A10^ GCs (green) and *Pf*CSP-specific Igh^g2A10^ PrePCs (orange) over time. **E**. Experimental schematic to examine BM PCs after immunization **F**. Representative flow cytometry plot plots demonstrating the gating strategy used to identify *Pf*CSP-specific Igh^g2A10^ PCs in the BM. **G**. The number of *Pf*CSP-specific Igh^g2A10^ PC per femur pair over time **H**. Logos plots of LC-CDR3s showing AA changes in *Pf*CSP-specific Igh^g2A10^ PCs over time. **I**. Frequency of J gene usage and occurrence of L114X and/or Q106X mutations in BM PCs overtime.

As the BM is the site of residence of long-lived PCs, we tracked Igh^g2A10^ PCs in this organ after immunization using Blimp1-GFP reporter mice ^38^ to facilitate the identification of BM PCs (Figure 4E-F). Despite an approximately constant formation of PrePCs in the GC we did not observe the expected increase in the number of BM PCs. Rather the number of BM PCs remained static after day 14 (Figure 4G). To determine the identity of the cells that form the BM PC population we single cell sorted these cells and sequenced them at days 7, 10, 14 and 84 post-immunization. This analysis showed that early unmutated cells were largely replaced by later emigrants carrying mutations at both position LC_114 and LC_106, though replacement was not absolute as a small number of unmutated cells were still detected at day 84 (Figure 4H-I). Overall, these data are consistent with the bulk of the BM PC compartment emerging after the peak of the GC response when the overall affinity of GC B cell is likely to be highest.

## Discussion

Our single cell RNA-seq data are consistent with a model in which the early waxing GC favours the recycling and expansion of cells in the dark zone but the later waning GC favours PC formation. As such our data are consistent with a temporal switch model of PC formation ^17^. Nonetheless, in absolute terms PC formation may still be significant at the peak of the GC response due to the size of the GC which may explain why others have reported a steadier output of cells from the GC ^18^.

Regardless it is likely that early arrivals in the BM are replaced by later cells due to the natural rate of decay^19,20^. Collectively these processes appear to be sufficient to result in a high affinity BM PC compartment even without the need for affinity-dependent selection of PrePCs in the GC. Conversely, our data would appear to contradict models in which high affinity cells efficiently obtain T cell help and then differentiate to become PCs ^15^. However previous work suggested that high affinity antigen-BCR affinity interactions may result in antigen not being available for processing and presentation on MHC Class II ^39,40^. Thus, high BCR affinity may not automatically translate to a greater ability to obtain T cell help. An excessive diversion of high affinity cells away from DZ recycling in the GC reaction may also be detrimental to the overall process of affinity maturation, thus the stochastic formation of PCs in the waning GC may be the most efficient strategy for generating protective immune responses.

## Acknowledgments

We thank M. Devoy, H. Vohra, and C. Gillespie of the Flow Cytometry Facility at the John Curtin School of Medical Research for assistance with flow cytometry. We further thank the Maxim Nekrasov and Peter Milburn for of the Biomedical Resource Facility at the John Curtin School of Medical Research for assistance with single cell RNA seq.

## Author Contributions

Conceptualization: HJS

Methodology: HJS, BP, ML, CD, DC, XG, HGK

Investigation: HJS, ML, CD, DC, XG, HGK

Visualization: HJS, BP

Formal Analysis: HJS, BP, TN

Resources: RAS, JT, AHI

Funding acquisition: IAC

Project administration: IAC

Supervision: JT, AHI, TN, IAC

Writing – original draft: HJS, IAC

Writing – review & editing: HJS, AHI, TN, IAC

## References

1. Plotkin, S.A. (2020). Updates on immunologic correlates of vaccine-induced protection. Vaccine 38, 2250–2257. 10.1016/j.vaccine.2019.10.046.

2. Amanna, I.J., Carlson, N.E., and Slifka, M.K. (2007). Duration of humoral immunity to common viral and vaccine antigens. N Engl J Med 357, 1903–1915. 10.1056/NEJMoa066092.

3. Slifka, M.K., Matloubian, M., and Ahmed, R. (1995). Bone marrow is a major site of longterm antibody production after acute viral infection. J Virol 69, 1895–1902. 10.1128/JVI.69.3.1895-1902.1995.

4. Slifka, M.K., and Ahmed, R. (1996). Long-term antibody production is sustained by antibody-secreting cells in the bone marrow following acute viral infection. Ann N Y Acad Sci 797, 166–176. 10.1111/j.1749-6632.1996.tb52958.x.

5. Jerne, N.K. (1951). A study of avidity based on rabbit skin responses to diphtheria toxin-antitoxin mixtures. Acta Pathol Microbiol Scand Suppl (1926) 87, 1–183.

6. Eisen, H.N., and Siskind, G.W. (1964). Variations in Affinities of Antibodies during the Immune Response. Biochemistry 3, 996–1008. 10.1021/bi00895a027.

7. Nieuwenhuis, P., and Opstelten, D. (1984). Functional anatomy of germinal centers. Am J Anat 170, 421–435. 10.1002/aja.1001700315.

8. Victora, G.D., Schwickert, T.A., Fooksman, D.R., Kamphorst, A.O., Meyer-Hermann, M., Dustin, M.L., and Nussenzweig, M.C. (2010). Germinal center dynamics revealed by multiphoton microscopy with a photoactivatable fluorescent reporter. Cell 143, 592–605. 10.1016/j.cell.2010.10.032.

9. Allen, C.D., Okada, T., Tang, H.L., and Cyster, J.G. (2007). Imaging of germinal center selection events during affinity maturation. Science 315, 528–531. 10.1126/science.1136736.

10. Smith, K.G., Light, A., Nossal, G.J., and Tarlinton, D.M. (1997). The extent of affinity maturation differs between the memory and antibody-forming cell compartments in the primary immune response. EMBO J 16, 2996–3006. 10.1093/emboj/16.11.2996.

11. Viant, C., Weymar, G.H.J., Escolano, A., Chen, S., Hartweger, H., Cipolla, M., Gazumyan, A., and Nussenzweig, M.C. (2020). Antibody Affinity Shapes the Choice between Memory and Germinal Center B Cell Fates. Cell 183, 1298–1311 e1211. 10.1016/j.cell.2020.09.063.

12. Wong, R., Belk, J.A., Govero, J., Uhrlaub, J.L., Reinartz, D., Zhao, H., Errico, J.M., D’Souza, L., Ripperger, T.J., Nikolich-Zugich, J., et al. (2020). Affinity-Restricted Memory B Cells Dominate Recall Responses to Heterologous Flaviviruses. Immunity 53, 1078–1094 e1077. 10.1016/j.immuni.2020.09.001.

13. Krautler, N.J., Suan, D., Butt, D., Bourne, K., Hermes, J.R., Chan, T.D., Sundling, C., Kaplan, W., Schofield, P., Jackson, J., et al. (2017). Differentiation of germinal center B cells into plasma cells is initiated by high-affinity antigen and completed by Tfh cells. J Exp Med 214, 1259–1267. 10.1084/jem.20161533.

14. Inoue, T., Moran, I., Shinnakasu, R., Phan, T.G., and Kurosaki, T. (2018). Generation of memory B cells and their reactivation. Immunol Rev 283, 138–149. 10.1111/imr.12640.

15. Ise, W., Fujii, K., Shiroguchi, K., Ito, A., Kometani, K., Takeda, K., Kawakami, E., Yamashita, K., Suzuki, K., Okada, T., and Kurosaki, T. (2018). T Follicular Helper Cell-Germinal Center B Cell Interaction Strength Regulates Entry into Plasma Cell or Recycling Germinal Center Cell Fate. Immunity 48, 702–715 e704. 10.1016/j.immuni.2018.03.027.

16. Gitlin, A.D., Shulman, Z., and Nussenzweig, M.C. (2014). Clonal selection in the germinal centre by regulated proliferation and hypermutation. Nature 509, 637–640. 10.1038/nature13300.

17. Weisel, F.J., Zuccarino-Catania, G.V., Chikina, M., and Shlomchik, M.J. (2016). A Temporal Switch in the Germinal Center Determines Differential Output of Memory B and Plasma CellsImmunity 44, 116–130. 10.1016/j.immuni.2015.12.004.

18. Robinson, M.J., Dowling, M.R., Pitt, C., O’Donnell, K., Webster, R.H., Hill, D.L., Ding, Z., Dvorscek, A.R., Brodie, E.J., Hodgkin, P.D., et al. (2022). Long-lived plasma cells accumulate in the bone marrow at a constant rate from early in an immune response. Sci Immunol 7, eabm8389. 10.1126/sciimmunol.abm8389.

19. Koike, T., Fujii, K., Kometani, K., Butler, N.S., Funakoshi, K., Yari, S., Kikuta, J., Ishii, M., Kurosaki, T., and Ise, W. (2023). Progressive differentiation toward the long-lived plasma cell compartment in the bone marrow. J Exp Med 220. 10.1084/jem.20221717.

20. Xu, A.Q., Barbosa, R.R., and Calado, D.P. (2020). Genetic timestamping of plasma cells in vivo reveals tissue-specific homeostatic population turnover. Elife 9. 10.7554/eLife.59850.

21. McNamara, H.A., Idris, A.H., Sutton, H.J., Vistein, R., Flynn, B.J., Cai, Y., Wiehe, K., Lyke, K.E., Chatterjee, D., Kc, N., et al. (2020). Antibody Feedback Limits the Expansion of B Cell Responses to Malaria Vaccination but Drives Diversification of the Humoral Response. Cell Host Microbe 28, 572–585 e577. 10.1016/j.chom.2020.07.001.

22. Fisher, C.R., Sutton, H.J., Kaczmarski, J.A., McNamara, H.A., Clifton, B., Mitchell, J., Cai, Y., Dups, J.N., D’Arcy, N.J., Singh, M., et al. (2017). T-dependent B cell responses to Plasmodium induce antibodies that form a high-avidity multivalent complex with the circumsporozoite protein. PLoS Pathog 13, e1006469. 10.1371/journal.ppat.1006469.

23. Anker, R., Zavala, F., and Pollok, B.A. (1990). VH and VL region structure of antibodies that recognize the (NANP)3 dodecapeptide sequence in the circumsporozoite protein of Plasmodium falciparum. Eur J Immunol 20, 2757–2761. 10.1002/eji.1830201233.

24. Glaros, V., Rauschmeier, R., Artemov, A.V., Reinhardt, A., Ols, S., Emmanouilidi, A., Gustafsson, C., You, Y., Mirabello, C., Bjorklund, A.K., et al. (2021). Limited access to antigen drives generation of early B cell memory while restraining the plasmablast response. Immunity 54, 2005–2023 e2010. 10.1016/j.immuni.2021.08.017.

25. Chan, T.D., Gatto, D., Wood, K., Camidge, T., Basten, A., and Brink, R. (2009). Antigen affinity controls rapid T-dependent antibody production by driving the expansion rather than the differentiation or extrafollicular migration of early plasmablasts. J Immunol 183, 3139–3149. 10.4049/jimmunol.0901690.

26. Paus, D., Phan, T.G., Chan, T.D., Gardam, S., Basten, A., and Brink, R. (2006). Antigen recognition strength regulates the choice between extrafollicular plasma cell and germinal center B cell differentiation. J Exp Med 203, 1081–1091. 10.1084/jem.20060087.

27. Espinosa, D.A., Christensen, D., Munoz, C., Singh, S., Locke, E., Andersen, P., and Zavala, F. (2017). Robust antibody and CD8(+) T-cell responses induced by P. falciparum CSP adsorbed to cationic liposomal adjuvant CAF09 confer sterilizing immunity against experimental rodent malaria infection. NPJ Vaccines 2. 10.1038/s41541-017-0011-y.

28. Seder, R.A., Chang, L.J., Enama, M.E., Zephir, K.L., Sarwar, U.N., Gordon, I.J., Holman, L.A., James, E.R., Billingsley, P.F., Gunasekera, A., et al. (2013). Protection against malaria by intravenous immunization with a nonreplicating sporozoite vaccine. Science 341, 1359–1365. 10.1126/science.1241800.

29. Nussenzweig, R.S., Vanderberg, J., Most, H., and Orton, C. (1967). Protective immunity produced by the injection of x-irradiated sporozoites of plasmodium berghei. Nature 216, 160–162. 10.1038/216160a0.

30. Pham, P., Bransteitter, R., Petruska, J., and Goodman, M.F. (2003). Processive AID-catalysed cytosine deamination on single-stranded DNA simulates somatic hypermutation. Nature 424, 103–107. 10.1038/nature01760.

31. Stoeckius, M., Hafemeister, C., Stephenson, W., Houck-Loomis, B., Chattopadhyay, P.K., Swerdlow, H., Satija, R., and Smibert, P. (2017). Simultaneous epitope and transcriptome measurement in single cells. Nat Methods 14, 865–868. 10.1038/nmeth.4380.

32. Laidlaw, B.J., Schmidt, T.H., Green, J.A., Allen, C.D., Okada, T., and Cyster, J.G. (2017). The Eph-related tyrosine kinase ligand Ephrin-B1 marks germinal center and memory precursor B cells. J Exp Med 214, 639–649. 10.1084/jem.20161461.

33. Suan, D., Krautler, N.J., Maag, J.L.V., Butt, D., Bourne, K., Hermes, J.R., Avery, D.T., Young, C., Statham, A., Elliott, M., et al. (2017). CCR6 Defines Memory B Cell Precursors in Mouse and Human Germinal Centers, Revealing Light-Zone Location and Predominant Low Antigen Affinity. Immunity 47, 1142–1153 e1144. 10.1016/j.immuni.2017.11.022.

34. Perez-Mazliah, D., Gardner, P.J., Schweighoffer, E., McLaughlin, S., Hosking, C., Tumwine, I., Davis, R.S., Potocnik, A.J., Tybulewicz, V.L., and Langhorne, J. (2018). Plasmodium-specific atypical memory B cells are short-lived activated B cells. Elife 7. 10.7554/eLife.39800.

35. Sutton, H.J., Aye, R., Idris, A.H., Vistein, R., Nduati, E., Kai, O., Mwacharo, J., Li, X., Gao, X., Andrews, T.D., et al. (2021). Atypical B cells are part of an alternative lineage of B cells that participates in responses to vaccination and infection in humans. Cell Rep 34, 108684. 10.1016/j.celrep.2020.108684.

36. Kennedy, D.E., Okoreeh, M.K., Maienschein-Cline, M., Ai, J., Veselits, M., McLean, K.C., Dhungana, Y., Wang, H., Peng, J., Chi, H., et al. (2020). Novel specialized cell state and spatial compartments within the germinal center. Nat Immunol 21, 660–670. 10.1038/s41590-020-0660-2.

37. Chen, S.T., Oliveira, T.Y., Gazumyan, A., Cipolla, M., and Nussenzweig, M.C. (2023). B cell receptor signaling in germinal centers prolongs survival and primes B cells for selection. Immunity 56, 547–561 e547. 10.1016/j.immuni.2023.02.003.

38. Kallies, A., Hasbold, J., Tarlinton, D.M., Dietrich, W., Corcoran, L.M., Hodgkin, P.D., and Nutt, S.L. (2004). Plasma cell ontogeny defined by quantitative changes in blimp-1 expression. J Exp Med 200, 967–977. 10.1084/jem.20040973.

39. Batista, F.D., and Neuberger, M.S. (2000). B cells extract and present immobilized antigen: implications for affinity discrimination. EMBO J 19, 513–520. 10.1093/emboj/19.4.513.

40. Batista, F.D., and Neuberger, M.S. (1998). Affinity dependence of the B cell response to antigen: a threshold, a ceiling, and the importance of off-rate. Immunity 8, 751–759. 10.1016/s1074-7613(00)80580-4.

