## Supplementary Text and Figures for "Plasma cells are formed in waning germinal centers via an affinity-independent process"


Key Resources Table
Resource Availability
Experimental Model Details
Method Details
Quantification and Statistical Analysis
Supplementary Figures 1-4
Supplementary dataset 1-3
Supplementary References

#### Key Resources Table

| Name | Source | Identifier |
|---|---|---|
| **Datasets** | | |
| $Igh^{g2A10}$ B cell 10X scRNA-seq datasets | GEO | |
| **Antibodies** | | |
| TotalSeq™-C0301 anti-mouse Hashtag 1 Antibody | Biolegend | 155861 |
| TotalSeq™-C0301 anti-mouse Hashtag 2 Antibody | Biolegend | 155863 |
| TotalSeq™-C0301 anti-mouse Hashtag 3 Antibody | Biolegend | 155865 |
| TotalSeq™-C0301 anti-mouse Hashtag 4 Antibody | Biolegend | 155867 |
| TotalSeq™-C0301 anti-mouse Hashtag 5 Antibody | Biolegend | 155869 |
| TotalSeq™-C0557 anti-mouse CD38 Antibody | Biolegend | 102735 |
| TotalSeq™-C0197 anti-mouse CD69 Antibody | Biolegend | 104551 |
| TotalSeq™-C0228 anti-mouse CD183 (CXCR3) Antibody | Biolegend | 126545 |
| TotalSeq™-C0106 anti-mouse CD11c Antibody | Biolegend | 117361 |
| Brilliant Violet 510™ anti-mouse CD45.1 Antibody | Biolegend | 110741 |
| Brilliant Violet 605™ anti-mouse IgD Antibody | Biolegend | 405727 |
| Brilliant Violet 605™ anti-mouse/human CD45R/B220 Antibody | Biolegend | 103243 |
| Pacific Blue™ anti-mouse/human GL7 Antigen (T and B cell Activation Marker) Antibody | Biolegend | 144614 |
| PE/Cyanine7 anti-mouse CD138 (Syndecan-1) Antibody | Biolegend | 142514 |
| BUV395 Rat Anti-Mouse CD19 | BD | 563557 |
| IgM Monoclonal Antibody (II/41), APC-eFluor™ 780, eBioscience™ | ThermoFisher | 47-5790-82 |
| CD38 Monoclonal Antibody (90), Alexa Fluor™ 700, eBioscience™ | ThermoFisher | 56-0381-82 |
| PerCP/Cyanine5.5 anti-mouse/human CD11b Antibody | Biolegend | 101228 |
| PerCP/Cyanine5.5 anti-mouse Ly-6G/Ly-6C (Gr-1) Antibody | Biolegend | 108428 |
| PerCP/Cyanine5.5 anti-mouse CD3 Antibody | Biolegend | 100218 |
| FITC anti-mouse IgD Antibody | Biolegend | 405704 |
| **Single cell RNA-seq reagents** | | |

| | | |
|---|---|---|
| Chromium Next GEM chip G Single cell Ki | 10X Genomics | 1000127 |
| Chromium Next GEM Single Library and Gel Bead Kit | 10X Genomics | 1000128 |
| Chromium Single cell 5' Feature Barcode Kit | 10X Genomics | 1000256 |
| Chromium Single cell Library Construction Kit | 10X Genomics | 1000190 |
| Chromium Single Cell Mouse BCR Amplification Kit | 10X Genomics | 1000255 |
| Single Index Kit N Set A | 10X Genomics | 1000212 |
| Single Index Kit T Set A | 10X Genomics | 1000213 |
| **Light Chain BCR sequencing reagents** | | |
| Agencourt AMPure XP beads | Beckman Coulter | A63880 |
| Betaine 5M | Sigma-Aldrich | B300-1VL |
| CloneAmp HiFi PCR Premix | Takara | 639298 |
| ERCC Spike-In Mix | ThermoFisher | 4456740 |
| RNAse inhibitor | Clontech | 2313A |
| SuperScript II RT | Life Technologies | 18064-071 |
| Triton X-100 | Sigma-Aldrich | T9284 |
| dNTP | ThermoFisher | R0182 |
| *Taq* DNA polymerase recombinant | ThermoFisher | EP0404 |
| **Primers** | | |
| mVkappaF: 5`-TCGTCGGCAGCGTCAGATGTGTATAAGAGACAGGAYATTGTGMTSACMCARWCTMCA-3' | ThermoFisher | |
| mCkappaR: 5`-GTCTCGTGGGCTCGGAGATGTGTATAAGAGACAGACTGAGGCACCTCCAGATGTT-3' | ThermoFisher | |
| Oligo-dT: 5'-AAGCAGTGGTATCAACGCAGAGTACT30VN-3' | ThermoFisher | |
| TS primer : 5'-AAGCAGTGGTATCAACGCAGAGTACATrGrG+G-3' | Qiagen | |
| ISPCR: 5'-AAGCAGTGGTATCAACGCAGAGT-3' | ThermoFisher | |

#### Resource Availability

*Lead contact*

*Materials availability*

This study did not generate new unique reagents

*Data and code availability*

Single cell RNA-seq data is available at the NCBI BioProject with the submission number GSE227370

**Experimental Model Details**

*Mice*

C57BL/6NCrl were purchased from the Australian Phenomics Facility (Canberra, ACT, Australia). Blimp1$^{GFP/+}$ (C57BL/6(Prdm1$^{tm1Nutt}$; MGI: 3510704) mice were a kind gift from Carola Vinuesa (The Australian National University) [1]. Igh$^{g2A10}$ were generated and described by us previously [2]. Mice were bred and maintained under specific pathogen free conditions in individually ventilated cages at the Australian National University. All animal procedures were approved by the Animal Experimentation Ethics Committee of the Australian National University (Protocol numbers: A2016/17 and A2019/36). All mice were 5-8 weeks old at the commencement of experiments. Within each experiment, mice were both age matched. Female mice were used throughout the experiments.

*Parasites*

Mice were immunized IV with 5x$10^4$ Pb-PfCSP sporozoites crossed to either an mCherry or GFP background in order to easily identify infected mosquitoes[3]. Sporozoites were collected by harvesting the salivary glands of Anopheles stephensi mosquitoes and were irradiated (200k Rad) using a MultiRad 225 (Flaxitron) irradiator prior to injection.

**Method Details**

*Flow Cytometry and sorting*

Spleen and bone marrow of mice were collected and were prepared into single cell suspension for flow cytometric analysis and sorting. splenocytes were isolated by mechanically disrupting the spleens through a 70µm nylon mesh filter while bone marrow cells were flushed from femurs and tibias with FACs buffer in 27g syringes. Cells were incubated with 1µg/mL Streptavidin and 10 µg/mL TruStain fcX antibody diluted in FACS buffer for 30min on ice. Cells were then incubated in a staining cocktail containing fluorescent antibodies and tetramer probes (as well as oligo-bound TotalSeq-C antibodies (Biolegend) for 10x sorting experiments) for 30 mins in the dark at 4°C. ACK lysis buffer (Sigma) was used to lyse red blood cells from the single cell suspensions and then cells were washed twice with FACS buffer. 1% 7AAD was then added as a Live/Dead dye immediately prior sorting or analysis. Flow-cytometric data was collected on a BD Fortessa or X20 flow cytometer (Becton Dickinson) and analyzed using FlowJo software (FlowJo). A BD FACs Aria I or II (Becton Dickinson) machine was used for FACS sorting of cells. $(NANP)_9$ tetramer probes were prepared in house by mixing PE or APC conjugated streptavidin (Invitrogen) with biotinylated $(NANP)_9$ peptide in a 1:4 molar ratio.

*Adoptive cell transfer*

The number of probe$^+$ CD19$^+$ Igh$^{g2A10}$ B cells to transfer were quantified from donor spleens via flow cytometry. $2x10^4$ probe$^+$ CD19$^+$ Igh$^{g2A10}$ B cells were then transferred into each recipient mouse via IV injection. Mice were immunized 1-2 days after adoptive transfer of donor cells.

*Single Cell Light Chain Sequencing*

RNA was extracted and cDNA synthesized from sorted, single Igh$^{g2A10}$ B cells using a modified SMARTseq 2 protocol (Picelli et al., 2014). Cells were sorted into plates with wells containing 1ul of the cell lysis buffer, 0.5 ml dNTP mix (10 mM) and 0.5 ml of the oligo-dT primer at 5mM. The amount reagent used in the following reverse-transcription and PCR amplification step by half the original protocol while the concentration of the IS PCR primer was also further reduced to 50nM. The BCR light chain was then amplified from the cDNA using a 10-94 specific forward primer with kappa specific reverse primer with the following thermocycler settings: 1 cycle at 95°C for 5 minutes followed by 50 cycles at 95°C for 1 minute, 43°C for 1 minute and 72°C for 1.5 minutes then finally 1 cycle at 72°C for 5 minutes before holding at 4° to cool. Amplified light chain sequences were then sequenced via sanger sequencing. Sequences were then analyzed using IMGT Vquest *scRNA-seq*

Probe$^+$ IgD$^-$ Igh$^{g2A10}$ B cells stained with TotalSeq-C feature and hashtag antibodies were sorted into FACS buffer then spun down. Cells were resuspended and loaded into a Chromium (10x Genomics). Indexed V(D)J, Feature Barcode and GEX libraries of sorted samples were prepared according to the protocol for Single Indexed 10X Genomics V(D)J 5' kit for mice with Feature barcoding kit (10X Genomics). Libraries were then sequenced using on an Illumina NovaSeq6000

*Generation of g2A10 antibody variants*

Constructs containing minigenes for the germline heavy (isotype: IgG2A) and light chains of the 2A10 antibody in a pcDNA3.1+ backbone were ordered commercially (Biomatik). To generate intermediate and low affinity 2A10 light chain variants, as well as those containing the F114L and Q106X AA changes mutations were introduced using the QuikChange II site directed mutagenesis kit according to the manufacturers instructions (Agilent). Antibodies were generated by transfecting HEK293T cells grown in DMEM supplemented with Nutridoma-SP (Roche) with 15 μg of each of the heavy and light chain plasmids in 0.06mg/ml branched PEI in 120mM NaCl. 3 and 6 days following transfections supernatants were collected, concentrated over a 100kDa Ultra-15 centrifugal filter unit, Ultracell-100 membrane (Amicon). Antibody concentrations were determined by sandwich ELISA on coats plated with anti-mouse kappa (Southern Biotech) as capture antibodies and horseradish peroxidase conjugated anti-mouse IgG2A (KPL) as detection antibodies.

*ELISA*

Binding of 2A10 antibody variants was determined by ELISA. Nunc Maxisorp Plates (Nunc-Nucleon) were coated overnight with 1ug/ml streptavidin followed by binding of biotinylated $(NANP)_9$ peptide for 1 hour. After blocking with 1% BSA, serial dilutions of the antibodies were incubated on the plates for 1 hour and after washing, incubated with HRP conjugated anti-IgG2A antibodies (KPL).

*Analysis of PfCSP binding kinetics*

Igh$^{g2A10}$ germline and mutant Fabs were recombinantly expressed (Genscript) and their binding to recombinant PfCSP (Genscript) was assessed using surface plasmon resonance (SPR). An HC30M chip (Carterra, 4279) was conditioned using successive injections of 50 mM NaOH, 0.5 M NaCl and 10 mM Glycine pH 2.0, and then primed twice with 25 mM MES with 0.05% Tween-20 before activation of the chip's surface with a 1:1 mixture of 400 nM EDC (Pierce, PG82079) and 100 mM NHS (Pierce, 24510). A 0.625 μg/mL concentration of each Fab was prepared using sodium acetate pH 4.5 with 0.05% Tween, and each Fab was coupled to the chip's surface in duplicate immediately following chip activation. Excess binding sites were blocked with 1M ethanolamine pH 8.5. A three-fold dilution series of PfCSP (0.23 – 500 nM) was injected onto the chip in ascending concentration without regeneration between successive injections. HEPES-buffered saline with Tween-20 and EDTA (HBSTE) with 0.05% BSA was used as running buffer throughout the assay. Binding data was collected over a 10 minute association phase and 30 minute dissociation phase for each antibody-antigen interaction. Data were analyzed using Kinetics Software (Carterra) and curves which deviated from the software-predicted model of 1:1 antigen-antibody binding interaction were excluded from the analysis of binding kinetics. Dissociation rates ($k_d$), association rates ($k_a$) as well as dissociation constants ($K_D$) were calculated using the Kinetics Software.

#### Quantification and Statistical Analysis

*Single Cell RNA transcriptomic analysis*

Cell Ranger (10X Genomics) was used to process the V(D)J, Feature Barcode and GEX sequences and prepare them for downstream analysis. *Seurat* (version 3.1)[4] was used for graph-based clustering and visualizations of the gene expression data and demultiplexing. All functions described are from *Seurat* or the standard R package (version 3.60) using the default parameters unless otherwise stated. Day 7 and Day 21 samples were initially analysed separately using the following procedures. Cells that expressed fewer than 200 genes and genes that were expressed in fewer than 3 cells were excluded, along with cells that had greater than 5% mitochondrial genes. Gene expression was normalized for the mRNA assay using the NormalizeData function. Centered Log-Ratio normalization was used for the CITE-seq assay. The 2000 most variable genes for each sample were identified using FindVariableFeatures. Expression of all genes was scaled using ScaleData to linearly regress out sources of variation and Principal Components Analysis (PCA) on the variable genes identified above was then run with RunPCA. Louvain clustering was conducted on both D7 and D21 samples via the FindNeighbours and FindCluster functions to identify clusters of Non-B cells, which were excluded from further analyses. The remaining cells in each sample were then normalized and scaled again as above. Demultiplexing of hash-tagged samples was done using the HTODemux. Samples were then combined using FindIntegrationAnchors and then Intergratedata to create a combined dataset. The combined dataset was then scaled again as above and a PCA was run. FindNeighbours and FindClusters was used again to identify clusters. DEGs were identified using FindAllMarkers. The clustering was visualized with Uniform Manifold Approximation and Projection (UMAP) dimensionality reduction using RunUMAP and plotted using DimPlot with umap as the reduction. Log-normalized gene expression data was visualized using violin plots (VlnPlots) as well as onto -UMAP plots (FeaturePlot). Heatmaps were generated using DoHeatmap. Genesets for LZ, DZ [5], MBC[6] and PrePCs [7] were added to the seurat object via the addModule and visualised via UMAP. Pseudotime analysis was performed using *Monocle 3*

*10x VDJ sequence analysis*

AssignGenes.py and MakeDb.py from Change-O, part of the Immcantation portal were used to convert the 10X V(D)J output from CellRanger into an AIRR community standardized format to allow for further downstream analysis. Inferred germline V and J sequences were added with CreateGermline.py.

*Statistical analyses*

Details of specific statistical tests and experimental design are given in the relevant figure legends. Mouse experiments had 3-5 mice per group and were performed either in duplicate or triplicate. All data points are plotted from all replicate experiments. ANOVA analyses were performed in R version 4.2.2 (The R Foundation for Statistical Computing) on the pooled data from all replicate experiments using the *lmer* function. Where data was pooled from multiple experiments, each experiment was included as a blocking factor in the analysis. Where data are plotted on a log-scale data were log-transformed prior to analysis. Logistic regression analyses were performed using the *glm* function and included specific mutations as fixed factors and mouse as the random factor. Abbreviations for p values are as follows: $p < 0.05$ = *, $p < 0.01$ = **, $p < 0.001$ = ***, $p < 0.0001$ = ****; with only significant p values shown. No blinding or randomization was performed, however all readouts (ELISA, flow cytometry and sequencing) are objective readouts that are not subject to experimental bias.

*Lineage tree analysis*

To reconstruct a lineage tree for each mouse over all cell types, a maximum likelihood estimation was used based on the HLP19 substitution model for B-cell somatic hypermutation (SHM)[8]. HLP19 is based on GY94 [9] which is a codon-based model for the evolution of protein-coding DNA sequences which provides better fit and synonymous/nonsynonymous substitution rate estimates than nucleotide-based models. HLP19 in addition models known hotspot (WRC, GYW, WA, TW) and cold spot (SYC, GRS) motifs arising in B-cell SHM that violate assumptions in standard phylogenetic substitution models like GY94. Separate trees were estimated for HC and LC (J2). The mean tree depth, in units of mutations/site, was estimated from separate subtrees for each cell type.

#### Supplementary Figure 1

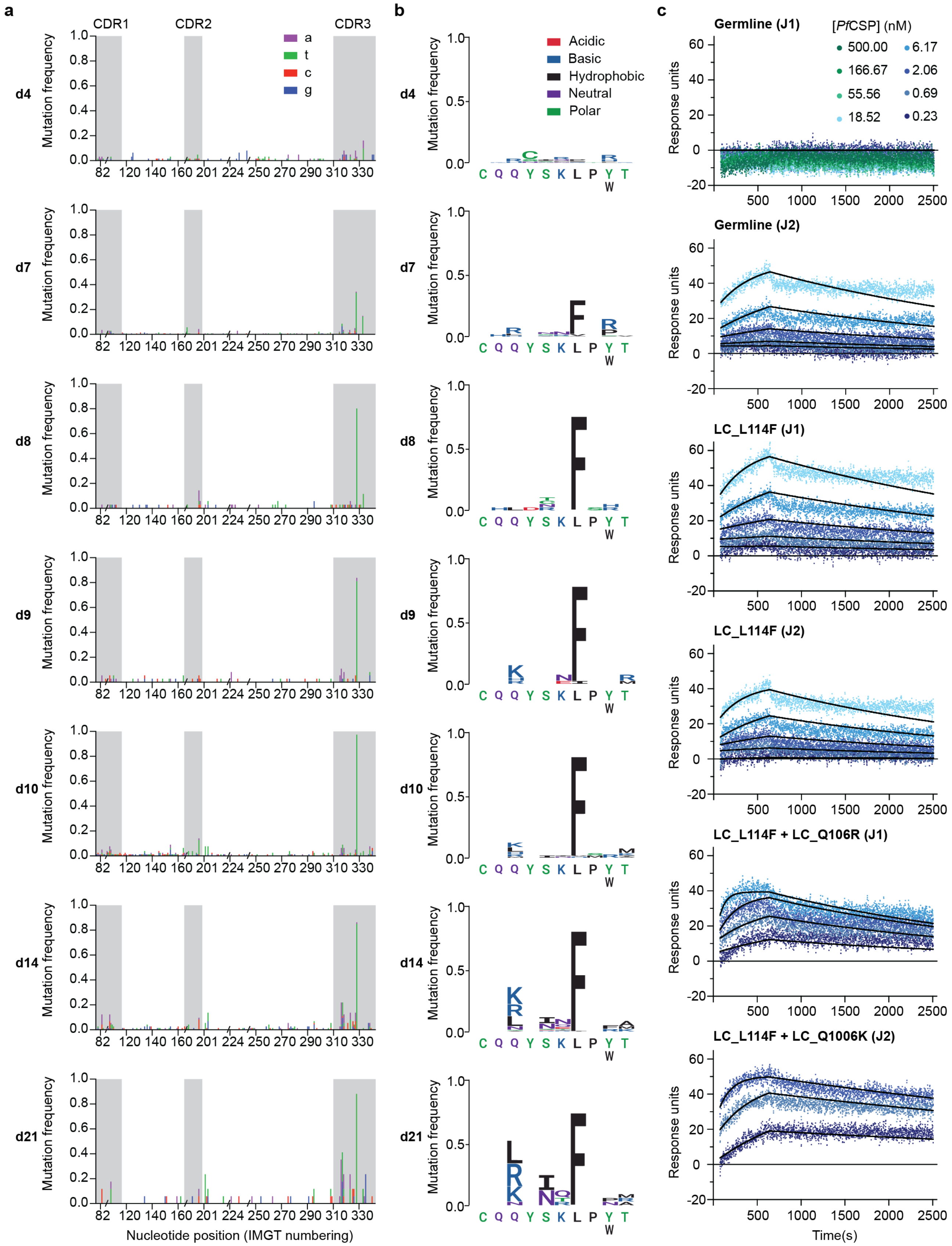


**Supplementary Figure 1: Mutation frequencies in Igh$^{g2A10}$ cells after immunization**. Mice were immunized and analyzed as described in figure 1G **A.** Manhattan plots showing the frequency of nucleotide changes in the light chain of among single $Igh^{g2A10}$ Cells 4-21 days post PbPf-SPZ immunizations. **B.** Logos plots of LC-CDR3s showing amino acid changes in *Pf*CSP-specific $Igh^{g2A10}$ B cells over time. **C.** Binding curves of $Igh^{g2A10}$ and mutant Fabs to *Pf*CSP**;** points show SPR sensorgram raw data while curves were generated from global 1:1 fit for binding of each Fab to PfCSP

### Supplementary Figure 2

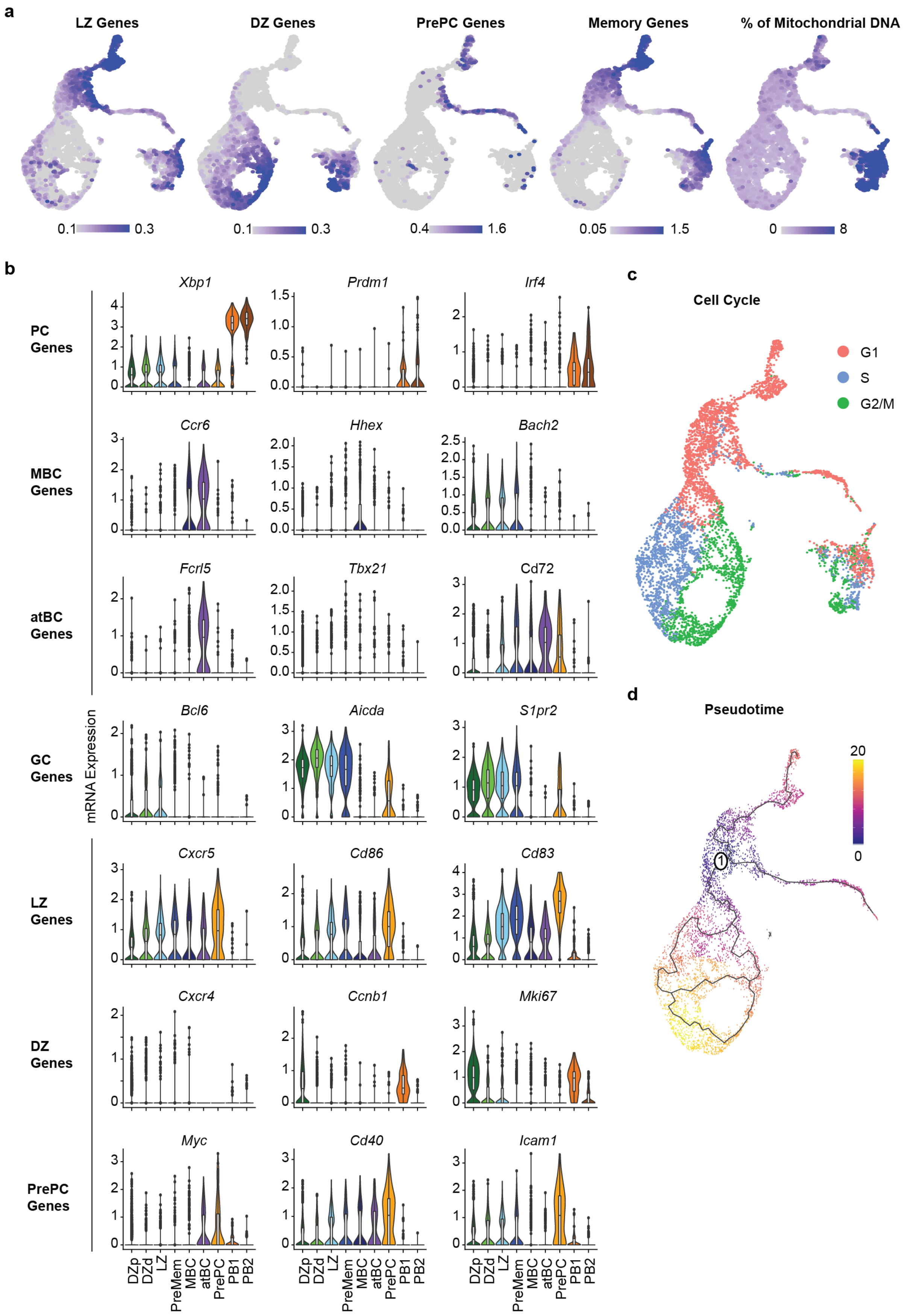

**Supplementary Figure 2: Gene set enrichment and candidate gene analysis to assign Igh$^{g2A10}$ B cell clusters.** Mice received Igh$^{g2A10}$ cells and were immunized with PbPf-SPZ and analyzed as described in Figure 2A. **A.** Enrichment scores for gene signatures distinguishing LZ, DZ PrePCs and MBCs as well as mitochondrial DNA expression projected onto UMAP plots. Colour was scaled for each gene signature **B.** Candidate gene analysis to assign Igh$^{g2A10}$ B cell clusters; violin and box plots showing the log-normalized expression mRNA expression of specific genes used to identify *Pf*CSP-specific Igh$^{g2A10}$ B cell clusters. **C.** Cell cycle stage for each cell projected onto a UMAP. **D.** Pseudotime analysis of B cells visualized using UMAP. Each point represents a cell and is coloured by cluster or progression along pseudotime. Pseudotime begins at "1", rooted on the LZ cluster.

### Supplementary Figure 3

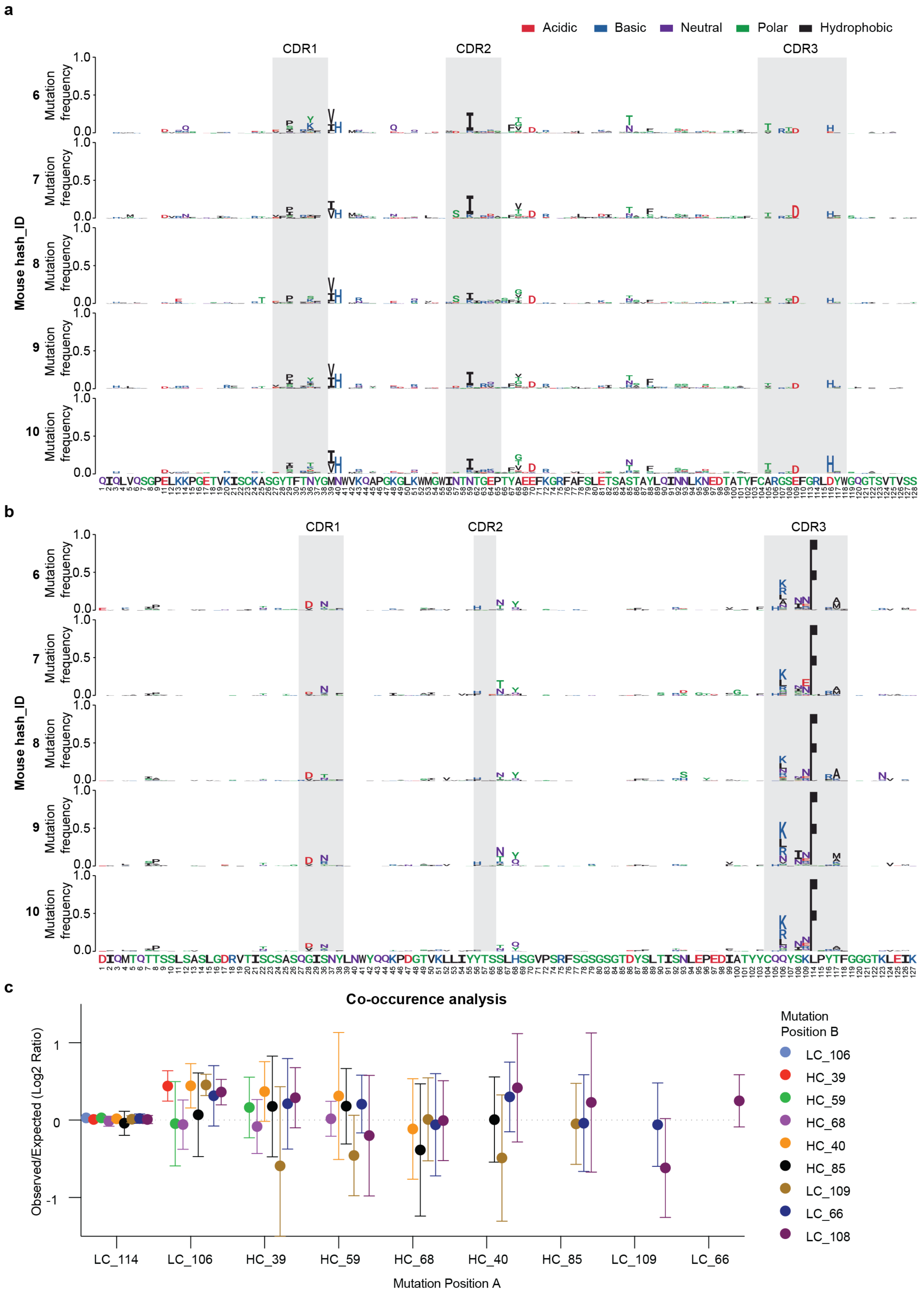

**Supplementary Figure 3 Location and co-occurrence of mutations in the Igh$^{g2A10}$ lineage.** Mice received Igh$^{g2A10}$ cells and were immunized with PbPf-SPZ and analyzed as described in Figure 2A. **A and B** Logos plots of the Heavy (A) and Light chain (B) sequences showing AA changes in *Pf*CSP-specific Igh$^{g2A10}$ B cells in individual mice 21 days post-PbPf-SPZ immunizations. **C.** Co-occurance of mutations at each of the top 10 most mutated locations in each mouse; data are expressed as the mean ± 95% CI log2 ratio of the observed co-occurrence compared to what would be expected by chance alone for each of the possible 45 pairs of mutated sites.

#### Supplementary Figure 4

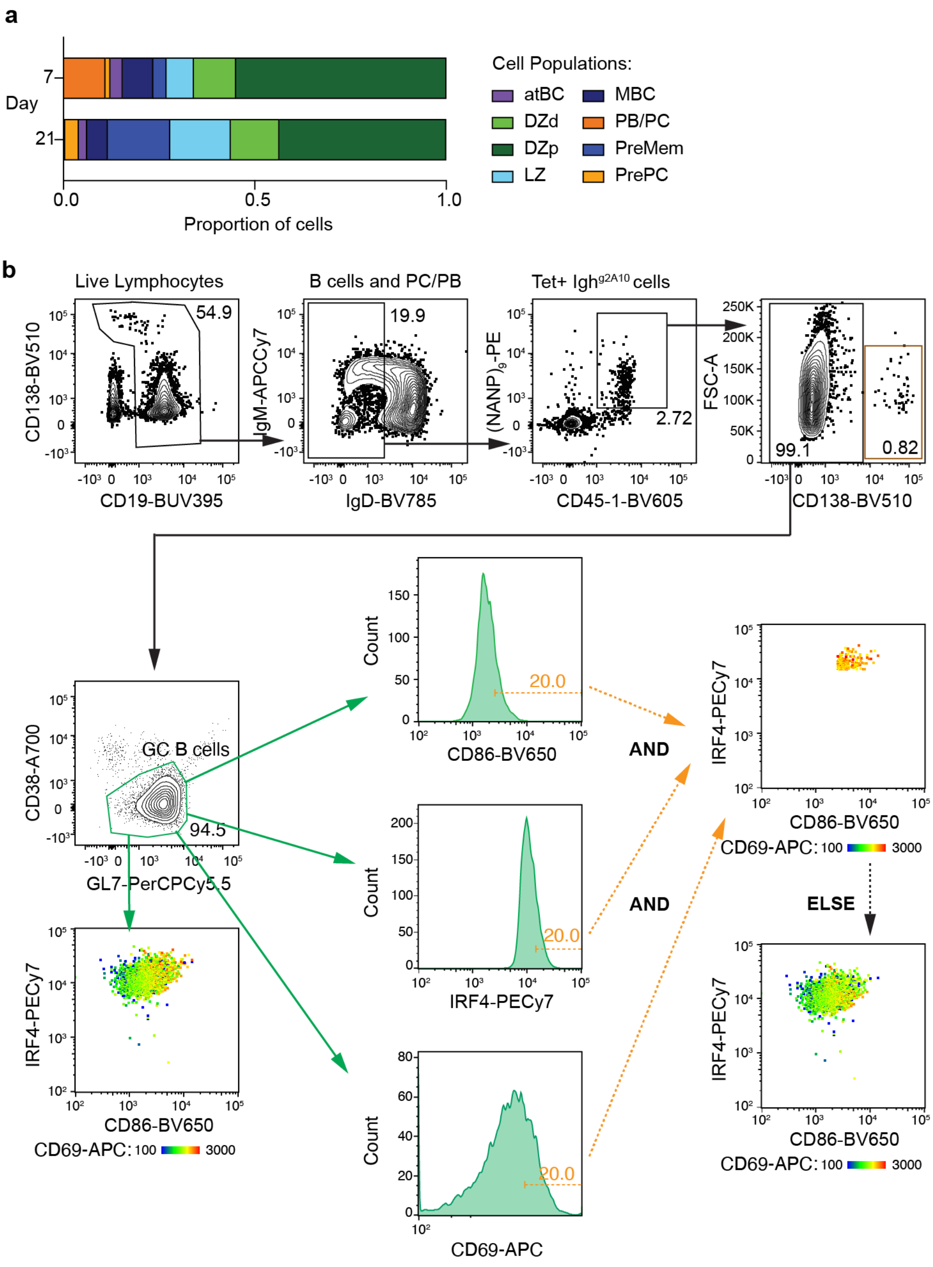




**Supplementary Figure 4: Quantitation of PrePCs in GCs A.** Mice received Igh$^{g2A10}$ cells and were immunized with PbPf-SPZ and analyzed as described in Figure 2A; proportion of *Pf*CSP-specific Igh$^{g2A10}$ B cells found in each cell population 7 or 21 days post-PbPf-SPZ immunizations from mice. **B.** Mice received Igh$^{g2A10}$ cells and were immunized with PbPf-SPZ and analyzed as described in Figure 4B; representative flow plots showing the gating strategy used to identify *Pf*CSP-specific Igh$^{g2A10}$ PrePCs, day 21 example shown.

**Supplementary Dataset 1**

Nucleotide and amino acid sequences from the sequencing of light chains from flow sorted $Igh^{g2A10}$ cells isolated from the spleen or bone marrow at different time points post immunization. Source data for Figure 1 D-H, Figure S1 and Figure 4 H-I.

**Supplementary Dataset 2**

Nucleotide sequences in AIRR format for each cell from the VDJ analysis determined from 10x Chromium sequencing. A total x cells from 10 mice (5 mice per timepoint) were included in the analysis after excluding any cells with incomplete or truncated sequences. A separate table with the corresponding amino acid sequences is also provided.

**Supplementary Dataset 3**

Lineage trees for each individual mouse at the day 21 timepoint based on a maximum likelihood estimation based on the HLP19 substitution model for B-cell somatic hypermutation (SHM). To present the key non-synonymous mutations within the context of the overall sequence evolution of the cells, the ancestral sequences of internal nodes were estimated and the first appearance of the key mutations are highlighted in the phylogenetic tree. The x-axis is scaled to mutations/base. Colors indicate the phenotype of each individual cell at the leaf nodes of the tree.

**Supplementary References**


1. Kallies, A., Hasbold, J., Tarlinton, D.M., Dietrich, W., Corcoran, L.M., Hodgkin, P.D., and Nutt, S.L. (2004). Plasma cell ontogeny defined by quantitative changes in blimp-1 expression. J Exp Med *200*, 967-977. 10.1084/jem.20040973.
2. McNamara, H.A., Idris, A.H., Sutton, H.J., Vistein, R., Flynn, B.J., Cai, Y., Wiehe, K., Lyke, K.E., Chatterjee, D., Kc, N., et al. (2020). Antibody Feedback Limits the Expansion of B Cell Responses to Malaria Vaccination but Drives Diversification of the Humoral Response. Cell Host Microbe *28*, 572-585 e577. 10.1016/j.chom.2020.07.001.
3. Espinosa, D.A., Christensen, D., Munoz, C., Singh, S., Locke, E., Andersen, P., and Zavala, F. (2017). Robust antibody and CD8(+) T-cell responses induced by P. falciparum CSP adsorbed to cationic liposomal adjuvant CAF09 confer sterilizing immunity against experimental rodent malaria infection. NPJ Vaccines *2*. 10.1038/s41541-017-0011-y.
4. Butler, A., Hoffman, P., Smibert, P., Papalexi, E., and Satija, R. (2018). Integrating single-cell transcriptomic data across different conditions, technologies, and species. Nat Biotechnol *36*, 411-420. 10.1038/nbt.4096.
5. Victora, G.D., Dominguez-Sola, D., Holmes, A.B., Deroubaix, S., Dalla-Favera, R., and Nussenzweig, M.C. (2012). Identification of human germinal center light and dark zone cells and their relationship to human B-cell lymphomas. Blood *120*, 2240-2248. 10.1182/blood-2012-03-415380.
6. Laidlaw, B.J., Schmidt, T.H., Green, J.A., Allen, C.D., Okada, T., and Cyster, J.G. (2017). The Eph-related tyrosine kinase ligand Ephrin-B1 marks germinal center and memory precursor B cells. J Exp Med *214*, 639-649. 10.1084/jem.20161461.
7. Ise, W., Fujii, K., Shiroguchi, K., Ito, A., Kometani, K., Takeda, K., Kawakami, E., Yamashita, K., Suzuki, K., Okada, T., and Kurosaki, T. (2018). T Follicular Helper Cell-Germinal Center B Cell Interaction Strength Regulates Entry into Plasma Cell or Recycling Germinal Center Cell Fate. Immunity *48*, 702-715 e704. 10.1016/j.immuni.2018.03.027.
8. Hoehn, K.B., Lunter, G., and Pybus, O.G. (2017). A Phylogenetic Codon Substitution Model for Antibody Lineages. Genetics *206*, 417-427. 10.1534/genetics.116.196303.
9. Goldman, N., and Yang, Z. (1994). A codon-based model of nucleotide substitution for protein-coding DNA sequences. Mol Biol Evol *11*, 725-736. 10.1093/oxfordjournals.molbev.a040153.
