## Supplementary Dataset 3 for "Plasma cells are formed in waning germinal centers via an affinity-independent process"

### Mouse 6: Heavy Chain

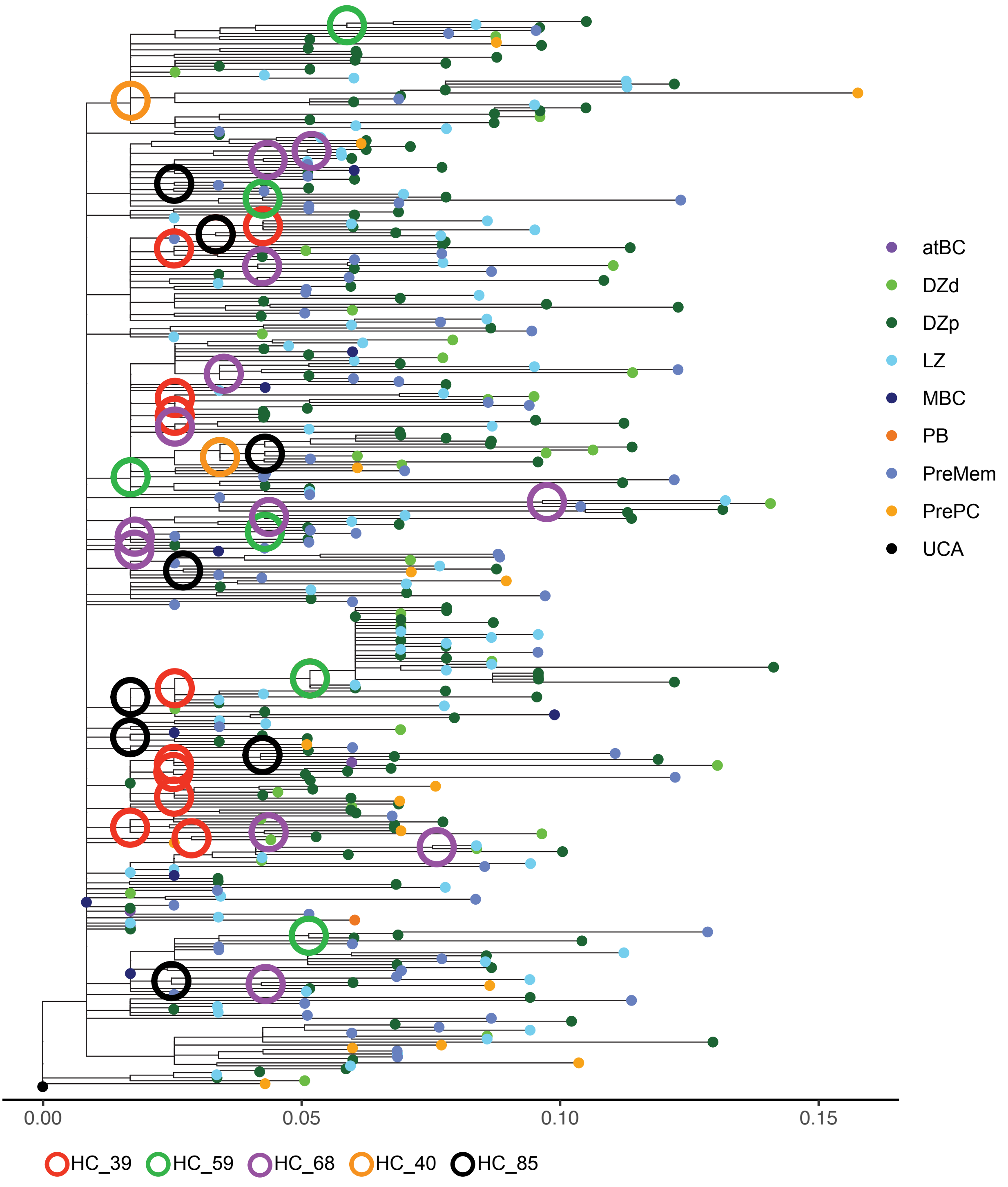

### Mouse 6: Light Chain (J2)

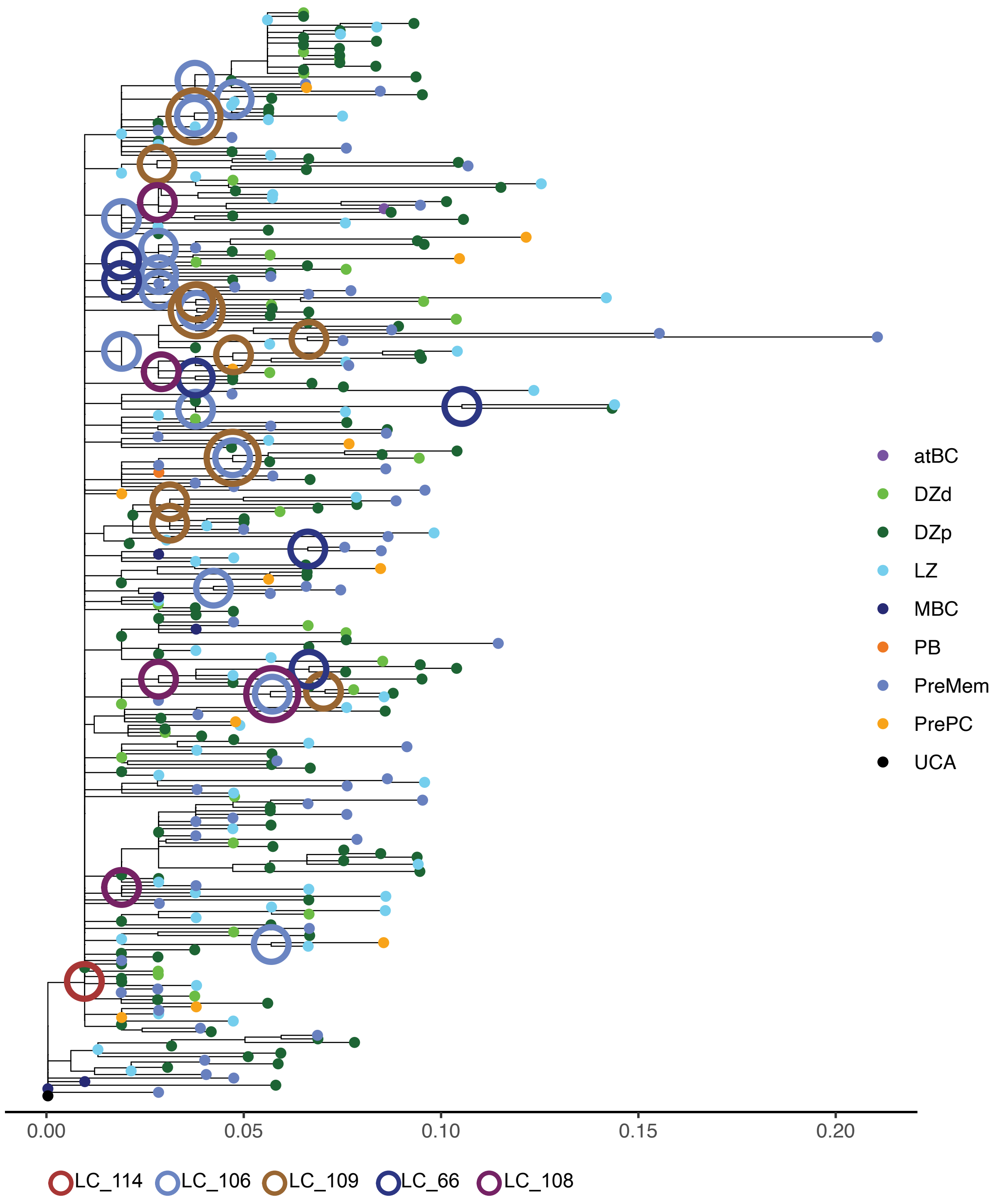

### Mouse 7: Heavy Chain

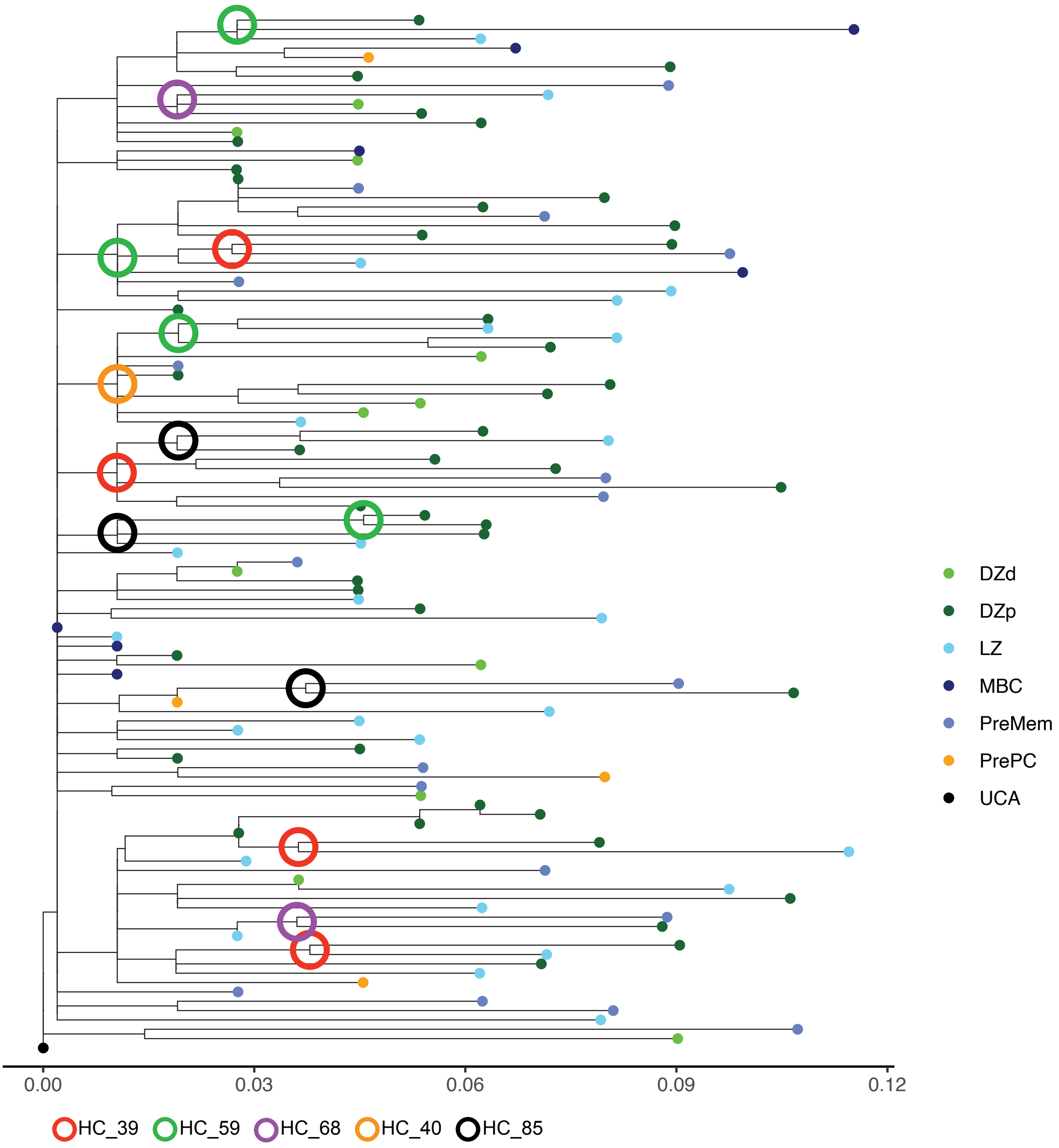

### Mouse 7: Light Chain (J2)

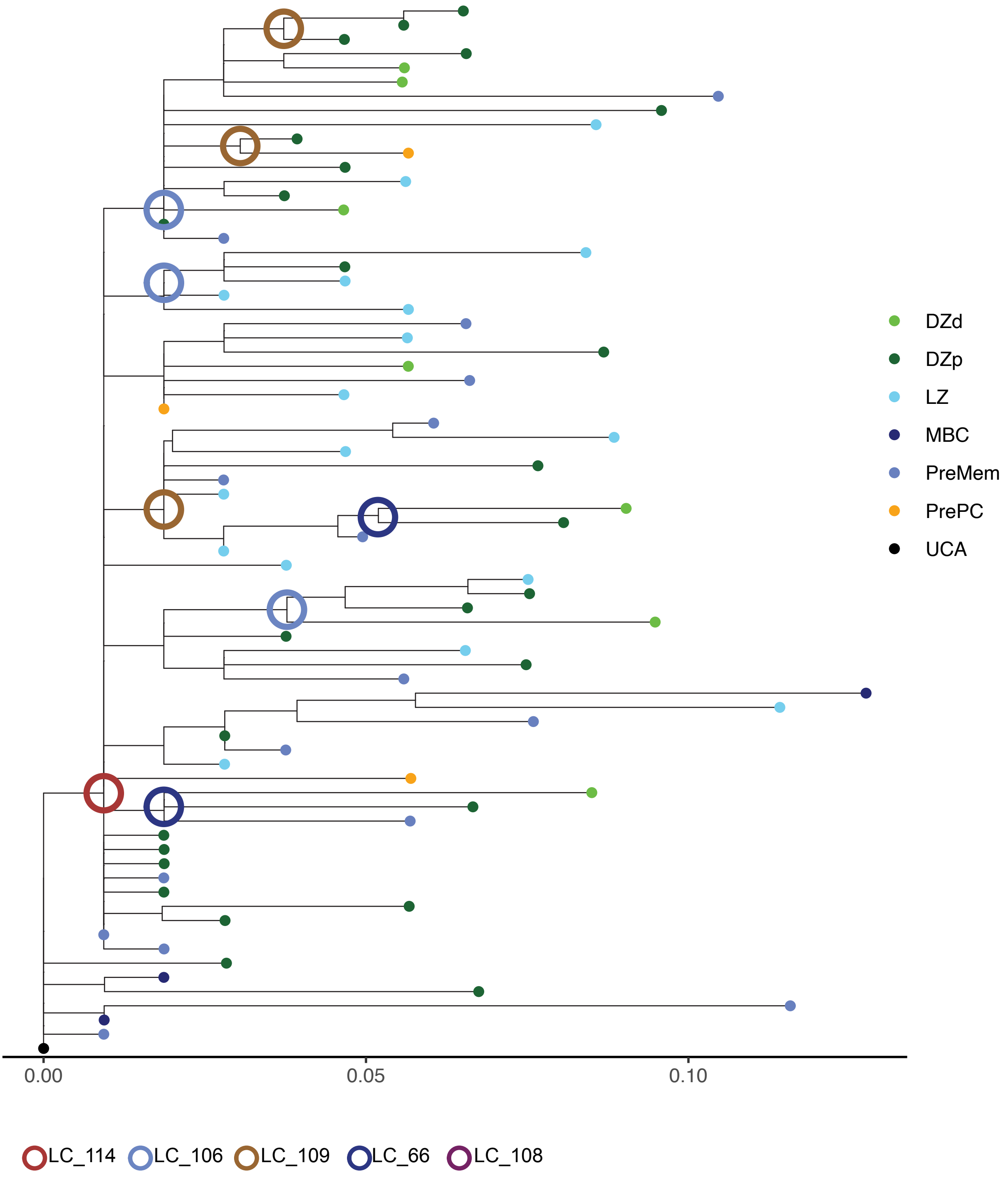

### Mouse 8: Heavy Chain

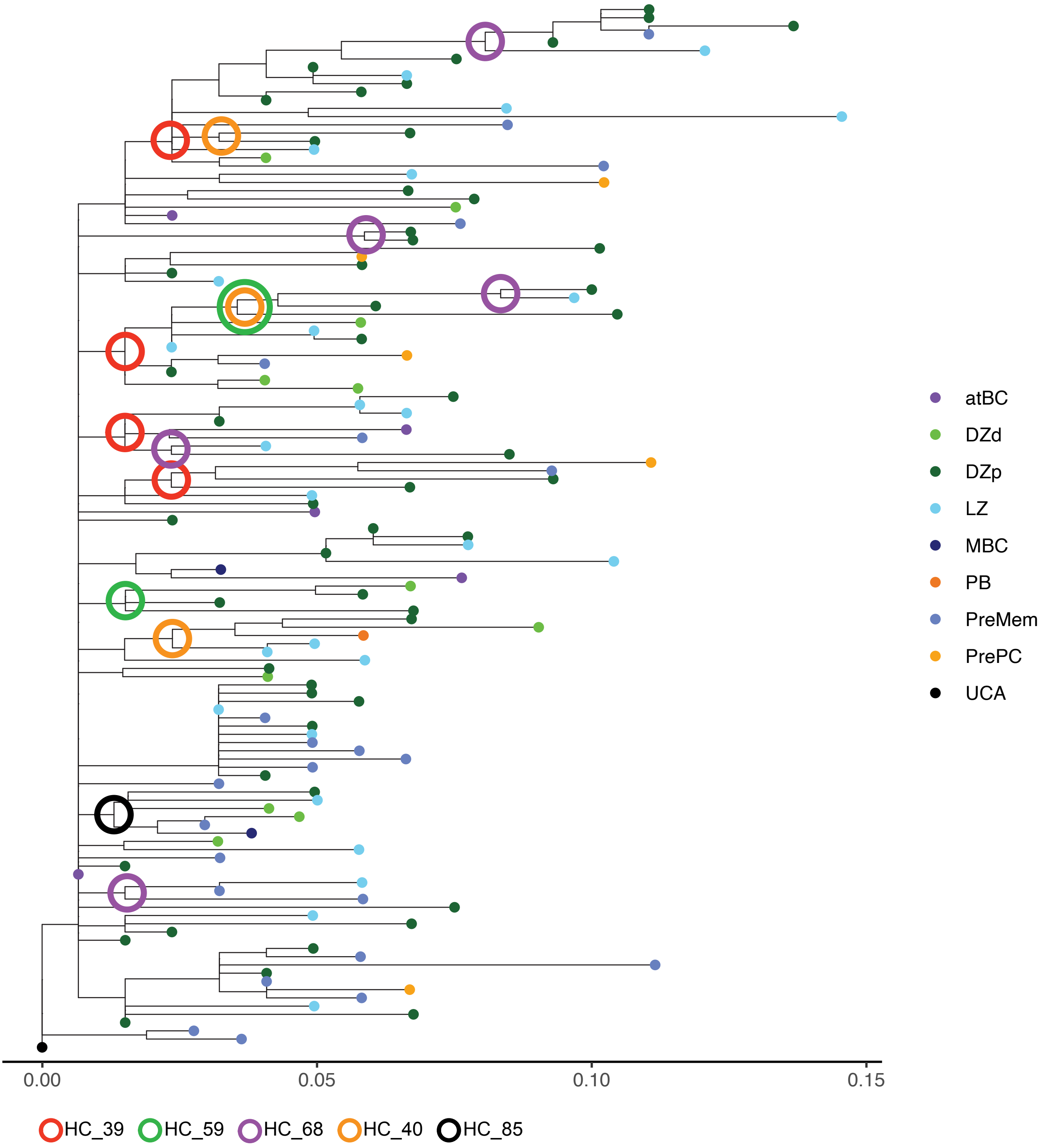

### Mouse 8: Light Chain (J2)

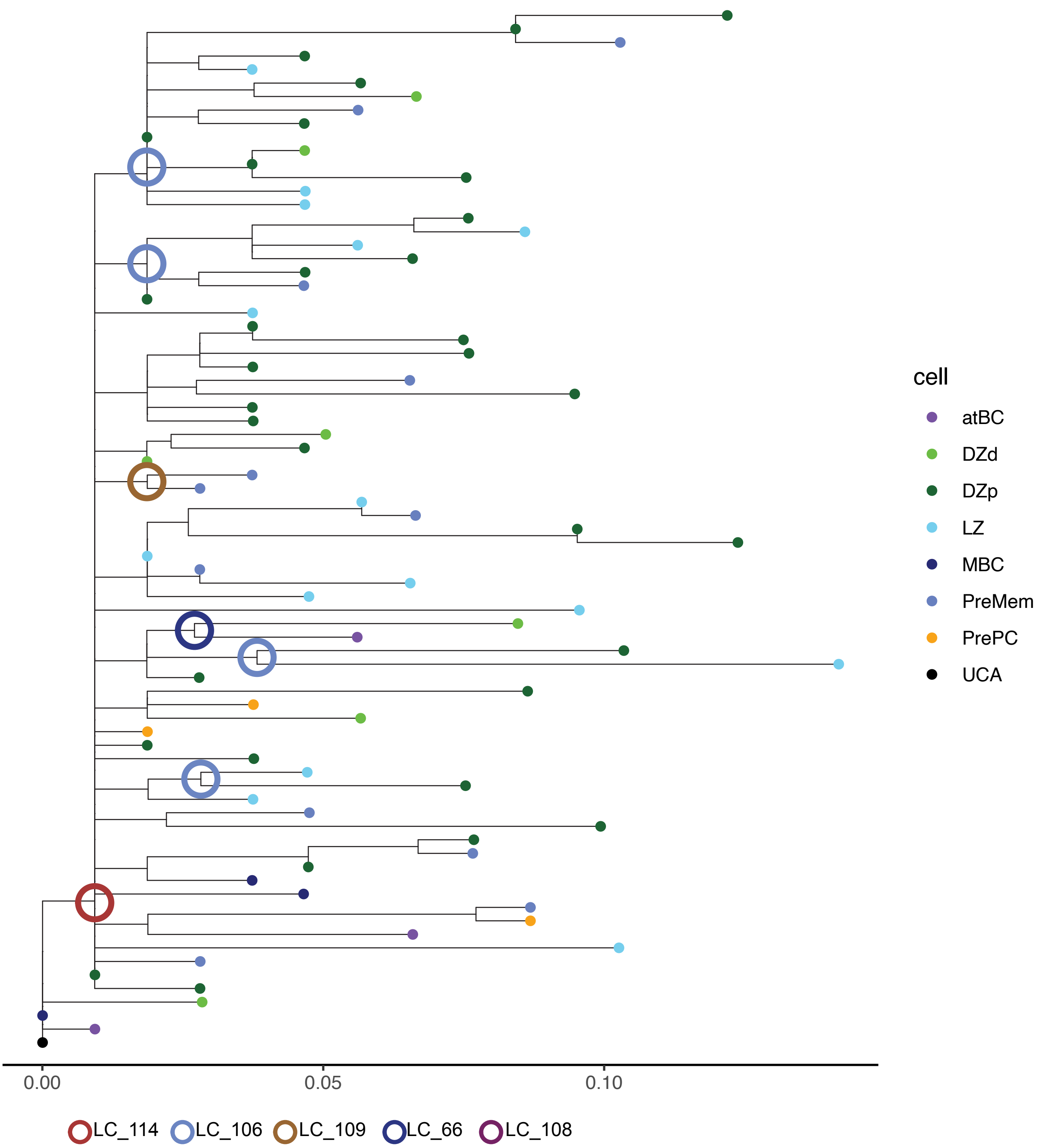

### Mouse 9: Heavy Chain

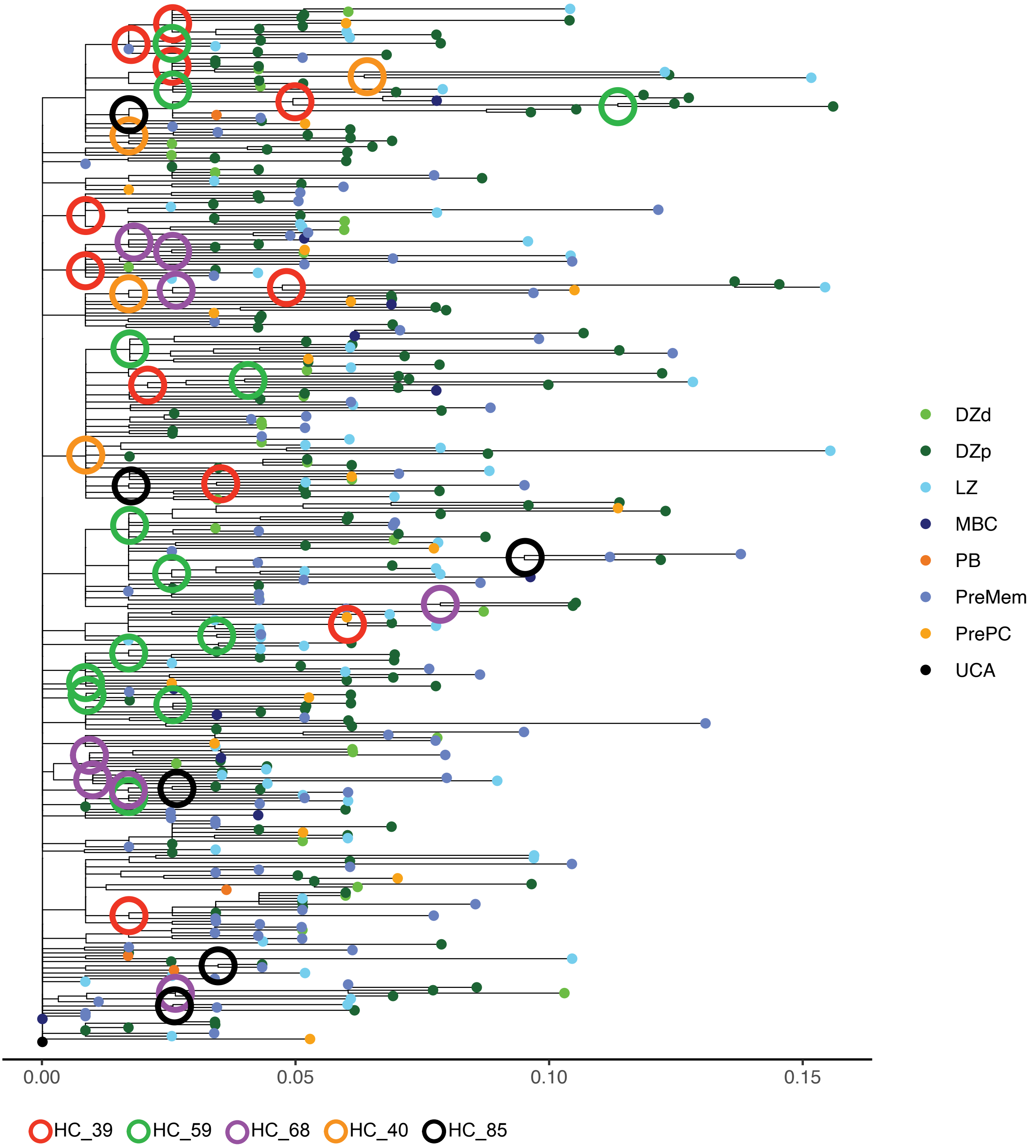

### Mouse 9: Light Chain (J2)

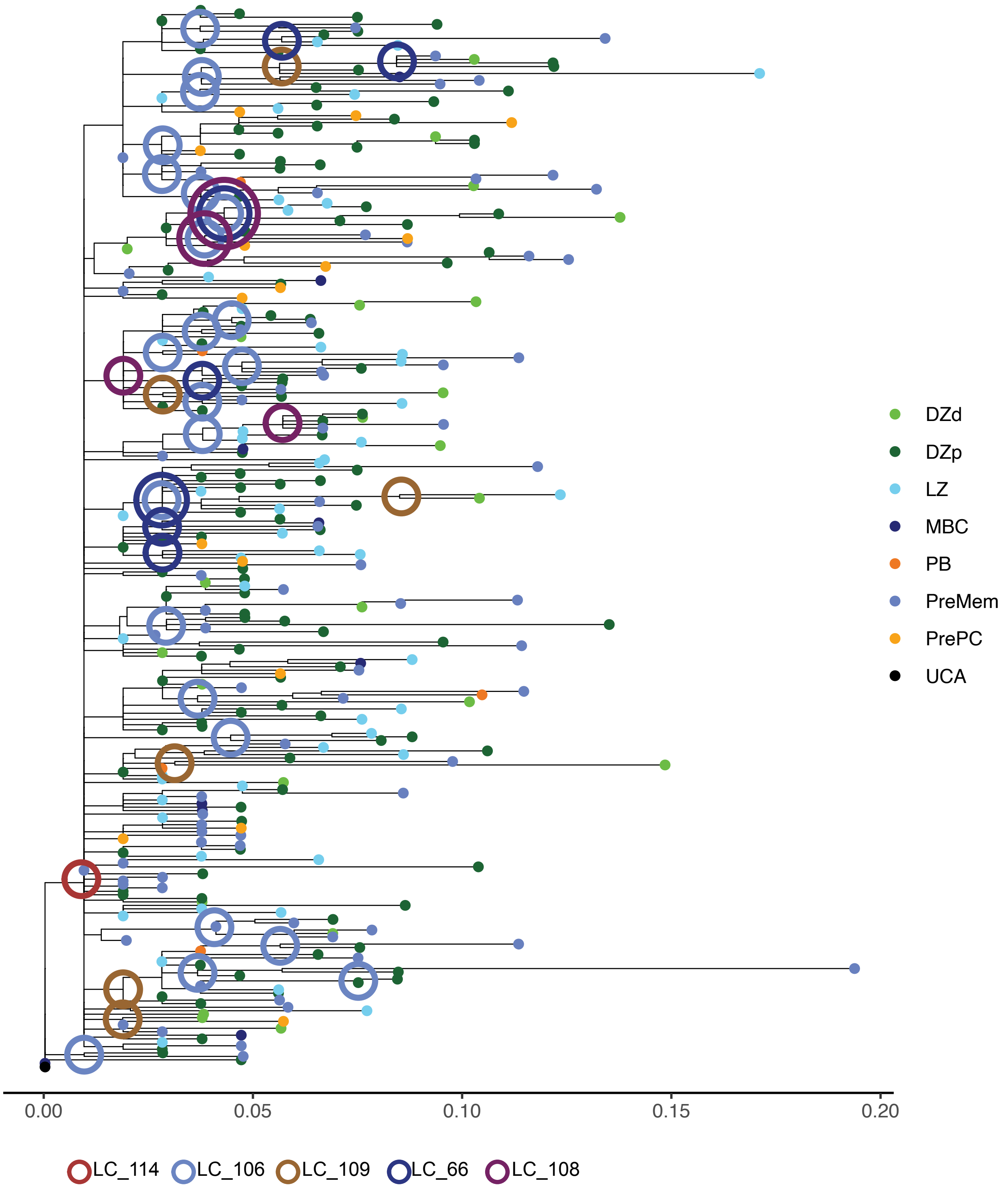

### Mouse 10: Heavy Chain

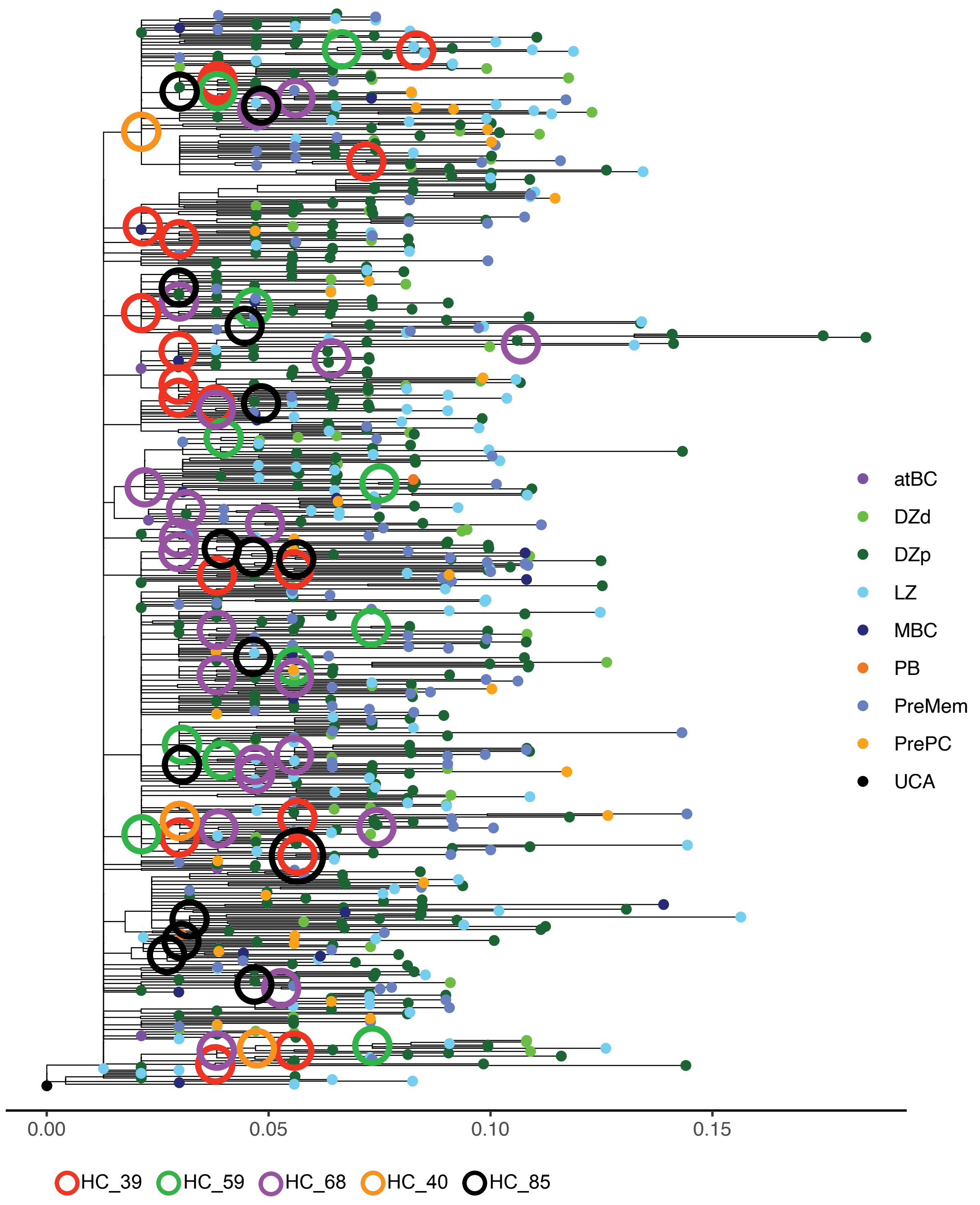

### Mouse 10: Light Chain (J2)

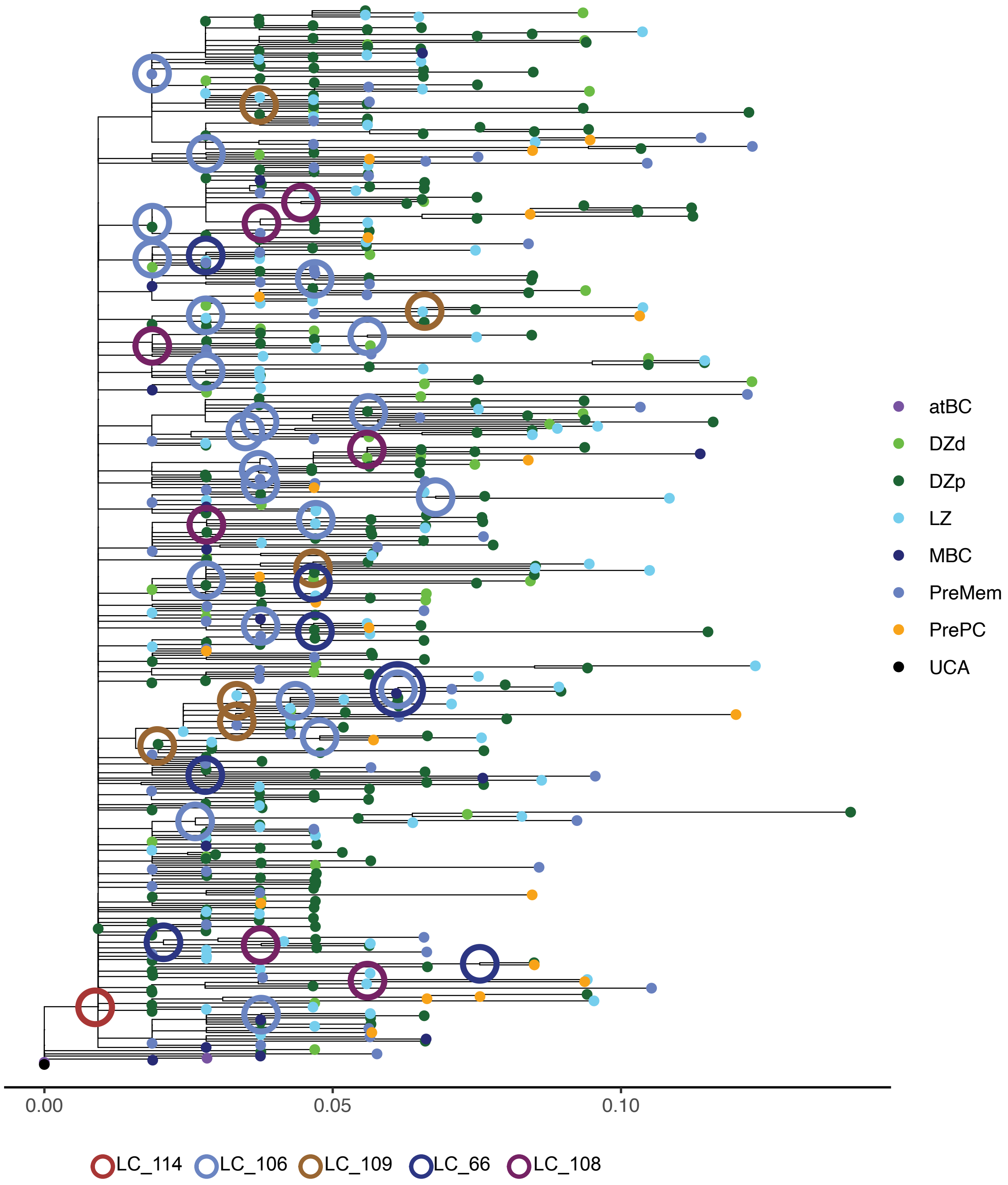
